# The SARS-CoV-2 Delta variant is poised to acquire complete resistance to wild-type spike vaccines

**DOI:** 10.1101/2021.08.22.457114

**Authors:** Yafei Liu, Noriko Arase, Jun-ichi Kishikawa, Mika Hirose, Songling Li, Asa Tada, Sumiko Matsuoka, Akemi Arakawa, Kanako Akamatsu, Chikako Ono, Hui Jin, Kazuki Kishida, Wataru Nakai, Masako Kohyama, Atsushi Nakagawa, Yoshiaki Yamagishi, Hironori Nakagami, Atsushi Kumanogoh, Yoshiharu Matsuura, Daron M. Standley, Takayuki Kato, Masato Okada, Manabu Fujimoto, Hisashi Arase

## Abstract

mRNA-based vaccines provide effective protection against most common SARS-CoV-2 variants. However, identifying likely breakthrough variants is critical for future vaccine development. Here, we found that the Delta variant completely escaped from anti-N-terminal domain (NTD) neutralizing antibodies, while increasing responsiveness to anti-NTD infectivity-enhancing antibodies. Although Pfizer-BioNTech BNT162b2-immune sera neutralized the Delta variant, when four common mutations were introduced into the receptor binding domain (RBD) of the Delta variant (Delta 4+), some BNT162b2-immune sera lost neutralizing activity and enhanced the infectivity. Unique mutations in the Delta NTD were involved in the enhanced infectivity by the BNT162b2-immune sera. Sera of mice immunized by Delta spike, but not wild-type spike, consistently neutralized the Delta 4+ variant without enhancing infectivity. Given the fact that a Delta variant with three similar RBD mutations has already emerged according to the GISAID database, it is necessary to develop vaccines that protect against such complete breakthrough variants.

## Introduction

Newly developed mRNA-based vaccines for SARS-CoV-2 have proven to be quite effective in preventing infection as well as severe COVID-19 (Jackson et al., 2020; Polack et al., 2020). However, new SARS-CoV-2 variants have repeatedly appeared and spread within the human population. Recent variants have acquired numerous mutations throughout the genome and are highly infectious compared to the original SARS-CoV-2. Although the spike protein used in currently approved mRNA-based vaccines consists of the original spike protein without mutations, these vaccines are nonetheless effective against variants of concern (VOC) (Collier et al., 2021; McCallum et al., 2021; Muik et al., 2021; Wang et al., 2021b). The receptor binding domain (RBD) of the spike protein binds to the host cell receptor ACE2, and the interaction mediates membrane fusion during SARS-CoV-2 infection (Hoffmann et al., 2020). Neutralizing antibodies against SARS-CoV-2 are mainly directed to the RBD and block the interaction between the RBD and ACE2. Most SARS-CoV-2 variants have acquired mutations in the neutralizing antibody epitopes of the RBD, resulting in escape from neutralizing antibodies (Cele et al., 2021; Collier et al., 2021; Davies et al., 2021; Madhi et al., 2021; Planas et al., 2021a; Tegally et al., 2021; Wang et al., 2021a). However, mutations in the RBD also tend to affect binding to ACE2. Therefore, there is a tradeoff in the evolution of the RBD between mutations that maintain ACE2 binding while escaping the recognition by neutralizing antibodies. In addition, mRNA vaccine-immune sera contain various neutralizing antibodies that recognize epitopes in different parts of the spike protein. It is an important to ascertain whether SARS-CoV-2 variants are likely to emerge that are completely resistant to immunity induced by the current mRNA-based vaccines. Vigilance against such resistant variants is essential for development of next-generation vaccines.

The SARS-CoV-2 Delta variant (B.1.617.2) is highly contagious and is rapidly spreading (Callaway, 2021). The neutralizing activity of sera from vaccinated individuals as well as convalescent COVID-19 patients decreases for the Delta variant compared to the wild-type (Liu et al., 2021a; Planas et al., 2021b). The Delta variant has several mutations in both the N-terminal domain (NTD) and RBD. The L452R and T478K mutations in the RBD of the Delta variant are also observed in other variants that are not as infectious as the Delta variant. Therefore, mutations in the RBD alone do not explain the high infectivity of the Delta variant. In contrast, among Delta mutations, several substitutions or deletions in the NTD—T19R, G142D, E156G, F157del and R158del—have not been observed in other major variants. This suggests that mutations in the NTD may play a key role in the high infectivity of the Delta variant. Although anti-RBD antibodies are thought to play a dominant role in vaccine-induced immunity against SARS-CoV-2 (Robbiani et al., 2020), neutralizing antibodies directed against the NTD are also important for SARS-CoV-2 neutralization (Chi et al., 2020; Li et al., 2021; Liu et al., 2020; Suryadevara et al., 2021; Voss et al., 2021). Moreover, we and others have recently demonstrated that antibodies against a specific site on the NTD can enhance the infectivity of SARS-CoV-2 by inducing the open form of the RBD (Li et al., 2021; Liu et al., 2021b). Therefore, it is important to elucidate the function of both the neutralizing and enhancing antibodies in order to understand the pathogenicity of the emerging SARS-CoV-2 variants. In this study, in order to understand the mechanism of the Delta variant’s high infectivity, we systematically examined Delta variant mutations in the NTD and RBD and suggest an evolutionary pathway by which the Delta variant could achieve complete escape from vaccine-induced immunity, which provides important information for the design of next-generation vaccines.

## Results

### Neutralizing activity of anti-NTD and anti-RBD monoclonal antibodies from COVID-19 patients against the Delta variant

In order to understand the mechanism underlying the increased infectivity of the SARS-CoV-2 Delta variant, we analyzed the binding of various types of anti-spike monoclonal antibodies obtained from COVID-19 patients to the Delta spike protein **(Figure 1A)**. Because these monoclonal antibodies were obtained from patients infected in mid-2020, at a time when the SARS-CoV-2 variants had not yet emerged, it is likely that they were elicited by the same wild-type spike protein as is used in current vaccines (Brouwer et al., 2020; Chi et al., 2020; Li et al., 2021; Robbiani et al., 2020; Suryadevara et al., 2021; Zost et al., 2020). Most neutralizing antibodies are directed against the RBD, and the Delta variant has two mutations in this domain, L452R and T478K. L452 has been reported to be an epitope for some, but not most, neutralizing antibodies (McCallum et al., 2021; Wang et al., 2021b). T478K is located in the ACE2 binding site and appears to be mainly involved in increased ACE2 binding affinity (Xu et al., 2021). In our analysis of various anti-RBD antibodies, we found that only a few of the neutralizing antibodies failed to recognize the Delta spike, while most anti-RBD neutralizing antibodies bound to Delta spike at levels comparable to wild-type spike **(Figure 1A)**.

**Figure 1.**
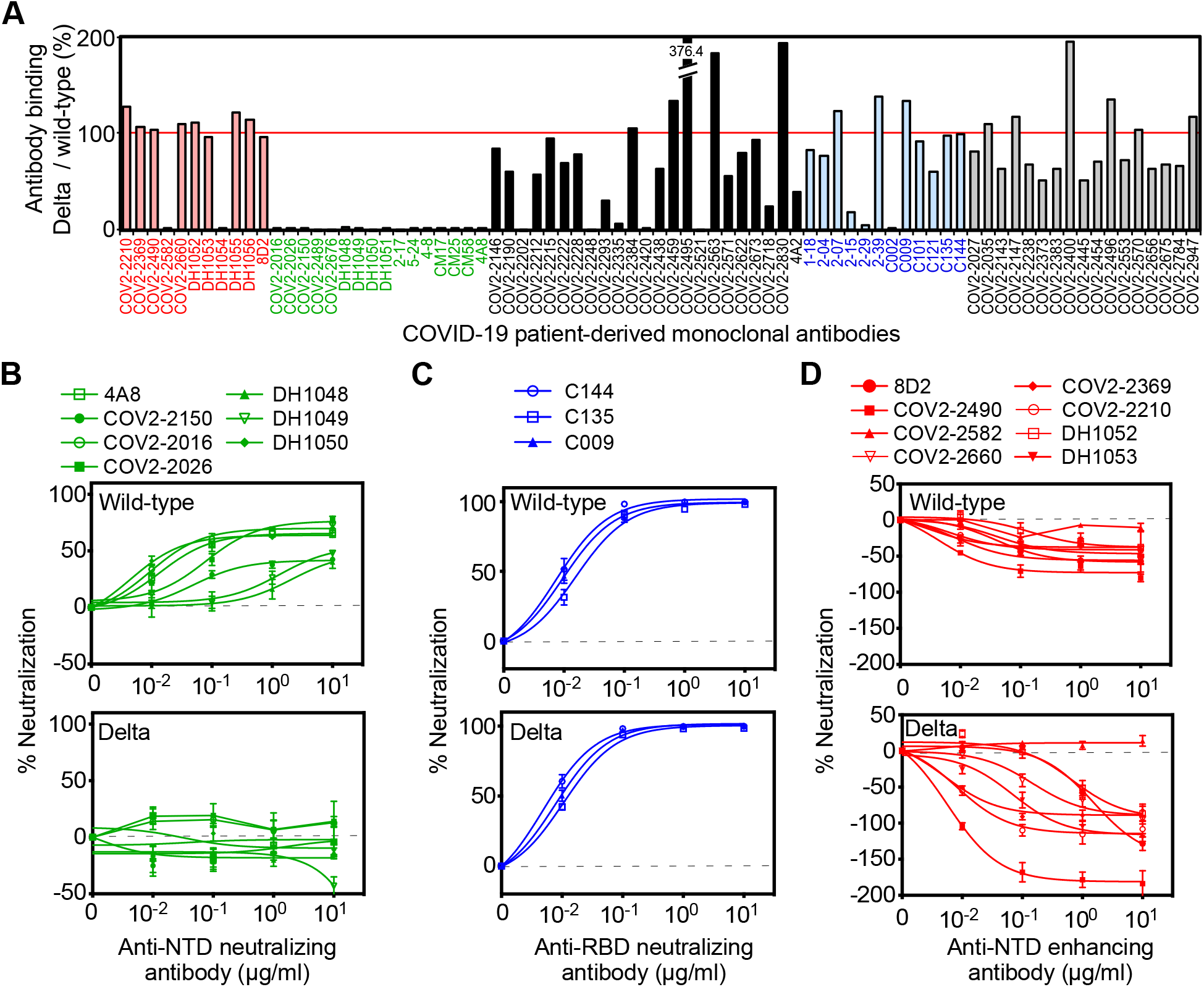
Neutralizing and enhancing effects against the wild-type and Delta spike pseudovirus by anti-spike monoclonal antibodies from COVID-19-patients. **(A)** The HEK293 cells transfected with the wild-type or the Delta spike were stained with anti-NTD enhancing antibodies (red), anti-NTD neutralizing antibodies (green), anti-NTD non-enhancing, non-neutralizing antibodies (black), anti-RBD neutralizing antibodies (blue) and anti-S2 antibodies (gray) (1 μg/ml). The stained cells were analyzed by flow cytometer. The relative mean fluorescence intensities (MFI) of antibodies binding to the Delta spike were compared with that for the wild-type spike. **(B-D)** The ACE2-expressing HEK293 cells were infected with the wild-type (upper) or the Delta (lower) pseudovirus in the presence of the anti-NTD neutralizing antibodies (B), anti-RBD neutralizing antibodies (C) and anti-NTD enhancing antibodies (D). A negative value for % neutralization indicates enhanced infectivity. The data from quadruplicates are presented as mean ± SEM. The representative data from three independent experiments are shown. See also Figure S1.

The Delta variant possesses several unique mutations in the NTD—T19R, G142D, E156G, F157del and R158del—suggesting the possibility that binding of some anti-NTD neutralizing antibodies elicited by wild-type spike could be disrupted. In addition to the 13 published anti-NTD neutralizing antibodies (Chi et al., 2020; Li et al., 2021; Liu et al., 2020; Suryadevara et al., 2021; Voss et al., 2021), we found that COV2-2016, COV2-2026 and COV2-2150 are also anti-NTD neutralizing antibodies for wild-type spike **(Figure 1B)**. We analyzed these 16 anti-NTD neutralizing antibodies, and found that none of the anti-NTD neutralizing antibodies could recognize Delta spike **(Figure 1A)**. In contrast, when we analyzed the binding of the anti-NTD infectivity-enhancing antibodoies (Li et al., 2021; Liu et al., 2021b), eight out of ten anti-NTD enhancing antibodies bound to Delta spike at levels comparable with wild-type spike **(Figure 1A)**. Some of the anti-NTD antibodies that were not well characterized as either neutralizing/enhancing antibodies showed partial or complete reduction in binding to Delta spike compared to wild-type spike, while others showed strong binding. The high frequency of reduced or enhanced recognition by anti-NTD antibodies against the Delta variant suggests that the antigenicity of the NTD has been greatly affected by mutations in the NTD.

Next, we analyzed the function of the enhancing and neutralizing antibodies on the Delta variants using pseudovirus bearing either the Delta spike protein (Delta pseudovirus) or wild-type spike (wild-type pseudovirus) **(Figure 1B–1D)**. The viral titer of each pseudovirus was checked by its infectivity to HEK293T cells transfected with ACE2 (**Figure S1**). Anti-RBD neutralizing antibodies that bound to the Delta spike completely neutralized the infection of either Delta or wild-type pseudovirus **(Figure 1C)**. All anti-NTD neutralizing antibodies we tested failed to recognize the Delta spike protein **(Figure 1A)**. As expected, these anti-NTD antibodies did not neutralize infection by the Delta pseudovirus, whereas they decreased the infectivity of the wild-type pseudovirus **(Figure 1B)**. The neutralizing efficiency of anti-NTD neutralizing antibodies against the wild-type pseudovirus was lower than that of anti-RBD neutralizing antibodies, as previously reported (Chi et al., 2020; Li et al., 2021; Liu et al., 2020; Suryadevara et al., 2021; Voss et al., 2021). Enhancing antibodies increase the infectivity of SARS-CoV-2 by inducing the open form of the RBD (Liu et al., 2021b). As described above, the recognition by most of the enhancing antibodies was well conserved in the Delta variant **(Figure 1A)**. When the effect of the enhancing antibodies was analyzed, the infectivity enhancement of the Delta pseudovirus by some of the enhancing antibodies was more than that of the wild-type pseudovirus **(Figure 1D)**. These data suggested that the Delta variant completely escaped from anti-NTD neutralizing antibodies while maintaining functional enhancing antibody epitopes. Because the enhancing antibodies decrease the effect of anti-RBD neutralizing antibodies (Li et al., 2021; Liu et al., 2021b), there is a possibility that the Delta variant maintains the infectivity in the presence of anti-RBD neutralizing antibodies as a result of enhancing antibodies.

### Neutralizing activity of BNT162b2-immune sera against Delta variants

We next analyzed the neutralizing activity of twenty sera from healthy individuals fully immunized with Pfizer-BioNTech BNT162b2 mRNA vaccine against the Delta pseudovirus **(Figure 2A)**. Although most of BNT162b2-immune sera completely blocked the infection of the Delta pseudovirus at high concentration, the neutralizing titer of BNT162b2-immune sera against Delta pseudovirus decreased significantly compared to wild-type pseudovirus **(Figure 2B)**, similar to a previous report (Liu et al., 2021a; Planas et al., 2021b). Because none of the anti-NTD neutralizing antibodies were effective against the Delta variant **(Figure 1A and 1B)**, it is likely that anti-RBD neutralizing antibodies play a major role in the neutralizing activity of BNT162b2-immune sera against the Delta variant.

**Figure 2.**
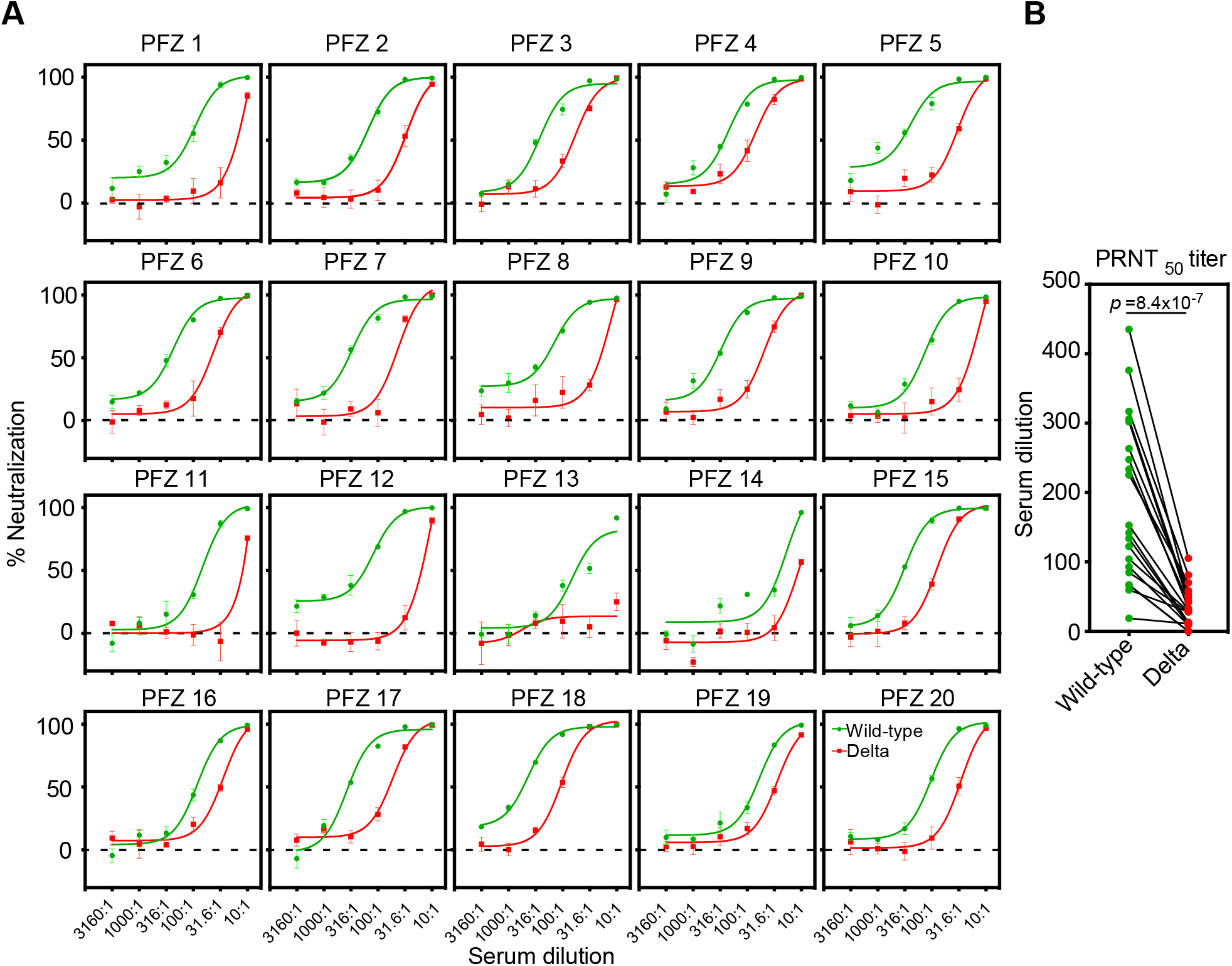
Neutralizing activity of BNT162b2-immune sera against the wild-type and the Delta pseudovirus. **(A)** Neutralizing activity of twenty BNT162b2-immune sera against the wild-type (green) and the Delta (red) pseudovirus. Data are mean ± SEM of technical quadruplicates. **(B)** PRNT50 titers of the BNT162b2-immune sera against the wild-type (green) and the Delta (red) pseudovirus are shown. *p* values determined by paired t-test were indicated. The representative data from three independent experiments are shown. See also Figure S1.

To elucidate the contribution of the NTD and RBD in the resistance of the BNT162b2-immune sera against the Delta variant, we generated chimeric spike proteins in which the NTD, RBD or S2 subunit was encoded by either the wild-type (W) or Delta (D) variant **(Figure 3A)**. Anti-NTD enhancing antibody, COV2-2490, binds to both the wild-type and Delta NTD, whereas anti-NTD neutralizing antibody, 4A8, binds to the wild-type NTD but not Delta NTD. Similarly, Anti-RBD neutralizing antibody, C144, binds to both the wild-type and Delta RBD, whereas anti-RBD neutralizing antibody, C002, binds to the wild-type RBD but not Delta RBD. As expected, C002 bound well to spike with the wild-type RBD (WWD or DWD) but weakly to spike with Delta RBD (DDD or WDD) **(Figure S2)**. Similarly, anti-NTD neutralizing antibody, 4A8, bound to spike with the wild-type NTD (WWD or WDD) but failed to bind to spike with the Delta NTD (DDD or DWD). COV2-2490 and C144 bound to all of the chimeric spike proteins. These data suggest that each domain of the chimeric spike proteins retains its original antigenicity.

**Figure 3.**
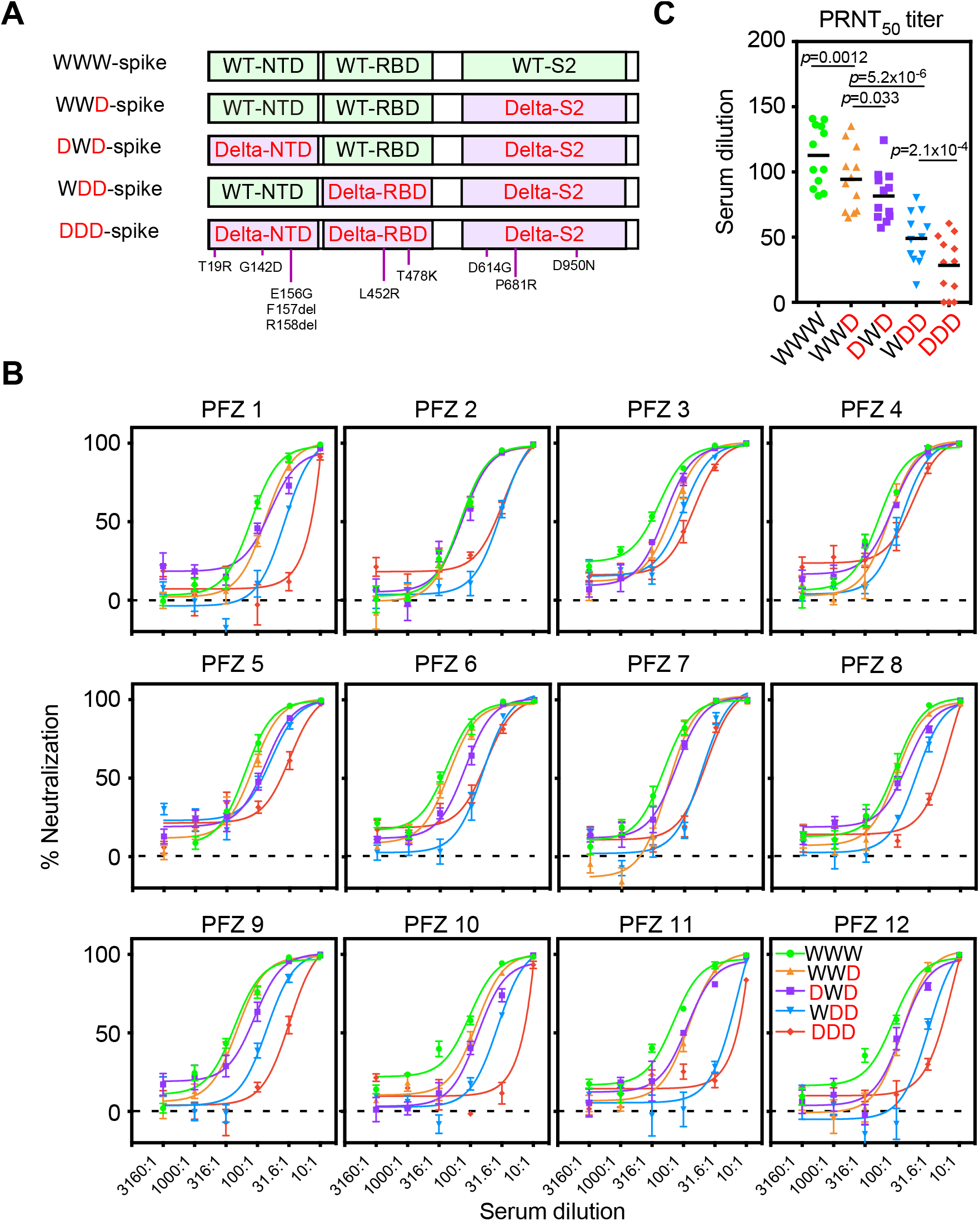
Neutralizing activity of BNT162b2-immune sera against the pseudovirus with chimeric spike protein of the wild-type and Delta variants. **(A)** The chimeric spike proteins between the wild-type (W) and Delta variant (D). Mutations of the Delta spike are indicated. **(B)** Neutralizing activity of BNT162b2-immune sera against the pseudoviruses with chimeric spike proteins. The data from quadruplicates are presented as mean ± SEM. **(C)** PRNT50 titers of BNT162b2-immune sera against the pseudoviruses with chimeric spike proteins. *p* values determined by paired t-test were indicated. The representative data from 2 independent experiments are shown. See also Figure S1 and S2.

We next generated pseudovirus containing these recombinant spike proteins and analyzed the effect of BNT162b2-immune sera. The neutralizing activity of the BNT162b2-immune sera against WWD pseudovirus decreased slightly compared to that of wild-type pseudovirus (WWW), suggesting that mutations in the S2 domain are involved in the resistance of the Delta variant **(Figure 3B and 3C)**. When infectivity of DWD pseudovirus, in which wild-type NTD was substituted to the Delta NTD, was compared with WWD pseudovirus, the neutralizing activity of BNT162b2-immune sera significantly decreased further. The neutralizing activity of the BNT162b2 immune sera was reduced against WDD pseudovirus, in which wild-type RBD was replaced by Delta RBD, compared to DWD pseudovirus. The neutralizing activity of the BNT162b2-immune sera decreased further against Delta pseudovirus (DDD). These data suggest that both NTD and RBD mutations in the Delta spike are involved in the resistance of the BNT162b2-immune sera against the Delta variant.

### Cryo-EM analysis of the Delta spike

All anti-NTD monoclonal neutralizing antibodies from COVID-19 patients failed to bind to Delta spike whereas most of the enhancing antibodies maintained reactivity to Delta spike (**Figure 1A**). Although there are several mutations in the NTD of Delta spike, known epitopes for anti-NTD neutralizing antibodies are conserved in the Delta variant. To evaluate the effect of mutations in the Delta variant on anti-NTD neutralizing antibody epitope structure, single particle cryo-EM analysis was employed. Data were analyzed by heterogenous refinement and *ab-initio* reconstruction followed by non-uniform refinement. As a result, a density map of the spike protein was obtained at 3.1 Å resolution **(Figure S3 and Table S1)**. To build an atomic model of the spike, we predicted the structures of the Delta variant NTD using AlphaFold2 (Jumper et al., 2021). The predicted NTD model of the Delta variant was used as an initial model for fitting into the obtained map. The statistics of the model of the Delta variant spike are summarized in **Table S1**. When the NTD models of Delta variant and wild-type spike were compared, the major epitope residues for the enhancing antibody—H64, W66, V213 and R214—were structurally well conserved (**Figure 4)**. In contrast, a large conformational change was observed in the residues of anti-NTD neutralizing antibody epitopes **(Figure 4)**. The maximum interatomic distance between the Delta variant and the wild-type was more than 9 Å **(Figure 4B)**. In the NTD of the Delta variant, the β strands containing four epitope residues—Y144, K147, K150 and W152—were shortened and shifted significantly compared to the wild-type **(Figure 4A)**. These structural changes were most likely caused by deletion of F157 and R158. As a result, these four residues were quite different from the wild type. R246 and W258 showed large changes compared to the wild-type **(Figure 4)**, and the loop connecting these two residues appeared to be highly flexible. These data suggest that dramatic changes in the structure of the anti-NTD neutralizing antibody epitope residues are responsible for the complete loss of reactivity to anti-NTD neutralizing antibodies against the Delta spike.

**Figure 4.**
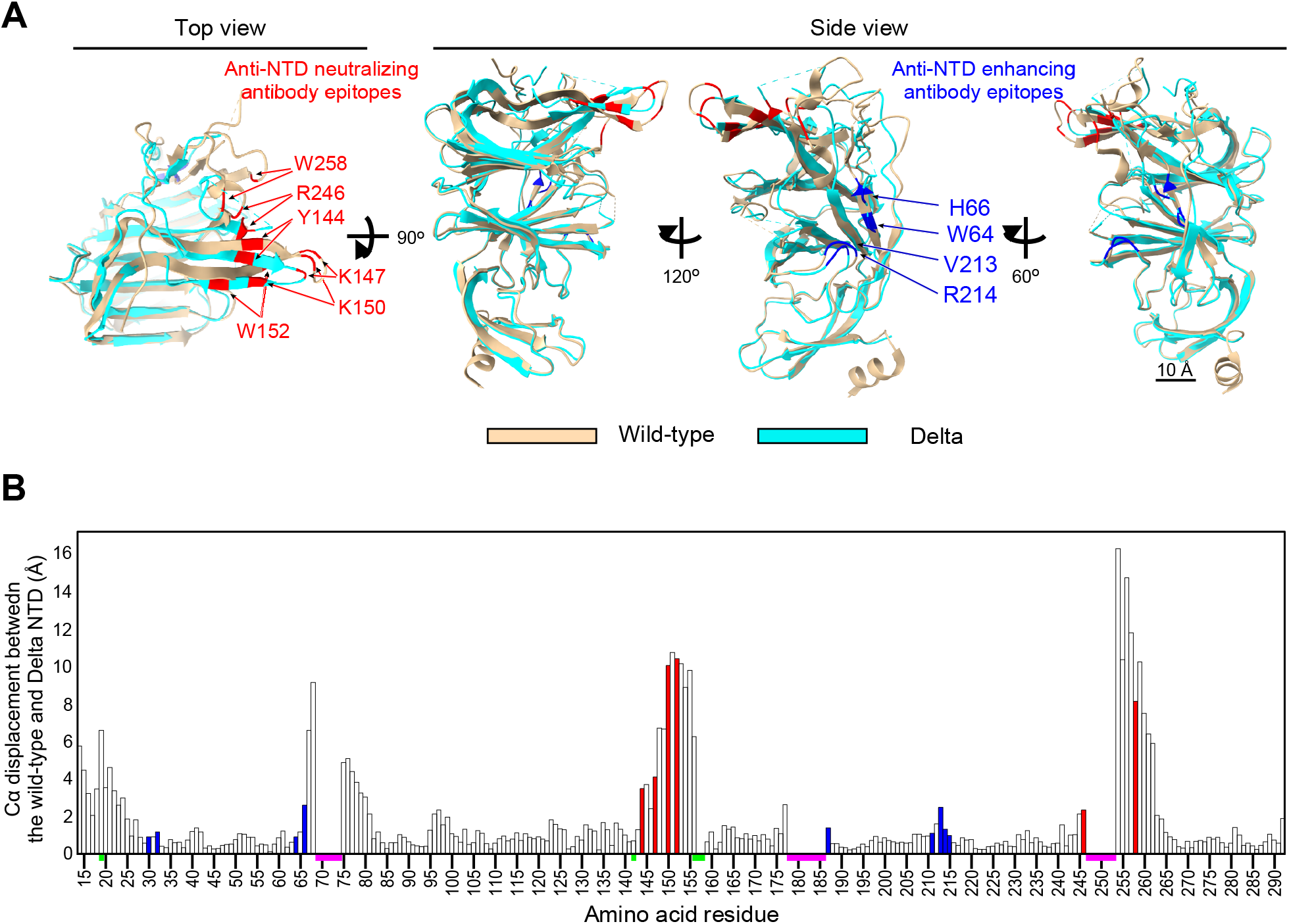
Cryo-EM analysis of the Delta NTD. **(A)** Structure of the Delta NTD (light blue) analyzed by the Cryo-EM were superimposed with the wild-type NTD (light brown, PDB: 7LY3). Major anti-NTD enhancing antibody epitopes (blue) and anti-NTD neutralizing antibody epitopes (red) were indicated in the figure. **(B)** Cα displacement between the wild-type and the Delta NTD was shown. The value was calculated by UCSF chimera. All known anti-NTD enhancing antibody epitopes (blue) and anti-NTD neutralizing antibody epitopes (red) were indicated. The regions where structures of wild-type or Delta NTD were not determined (magenta), and mutations in the Delta NTD (green) are indicated on the axis. See also Figure S3 and Table S1.

### Prediction of possible future mutations of the Delta variant

The Delta variant became completely resistant to anti-NTD neutralizing antibodies in the BNT162b2 immune serum by acquiring mutations in the NTD, and thus anti-RBD neutralizing antibodies seem to be mainly responsible for the neutralizing activity in the BNT162b2 immune sera **(Figure 1, Figure 2 and Figure 3)**. These results suggest that the Delta variant may acquire full resistance to BNT162b2 immune sera by acquiring additional mutations in the RBD that disrupt recognition of anti-RBD neutralizing antibodies. Indeed, a Delta variant that has acquired the K417N mutation in the RBD, known as AY.1 (Delta plus), has already emerged and its frequency in the general population is increasing (Gupta et al., 2021). To investigate the potential occurrence of additional mutations, we analyzed the additive effects of mutations acquired by the Delta variant in the GISAID database (**Figure S4**). The Delta variant has already acquired large numbers of additional mutations in the RBD, some of which occur in epitopes for anti-RBD neutralizing antibodies (Greaney et al., 2021a; Greaney et al., 2021b; Greaney et al., 2021c; Wang et al., 2021b; Weisblum et al., 2020). In addition to the K417N mutation, Delta variants with E484K, F490 or N501Y mutations—observed in the Alpha, Beta, Gamma and/or Lambda variants—are also increasing **(Figure 5A)**. Considering the very rapid increase in the population of people infected with the Delta variant, the Delta variant is likely to acquire further mutations in infected people, and those with further increased infectivity will be selected. Indeed, the Delta variant with multiple mutations in anti-RBD neutralizing antibody epitopes have already emerged according to the GISAID database **(Figure 5B)**. In particular, EPI_ISL_2958474 possesses three additional mutations in anti-RBD neutralizing antibody epitopes, although the NTD sequence is not identical to the representative Delta variant. Accordingly, we analyzed the effect of major mutations observed in SARS-CoV-2 variants on the RBD of the Delta variant (**Figure 5C**). Because the Delta variant contains the T478K mutation and neighboring residues may show similar effects, the S477N mutation was excluded. Accordingly, we introduced four mutations in the Delta spike (Delta 4+)—K417N, N439K, E484K and N501Y—and analyzed the effect of these mutations (**Figure 5D**).

**Figure 5.**
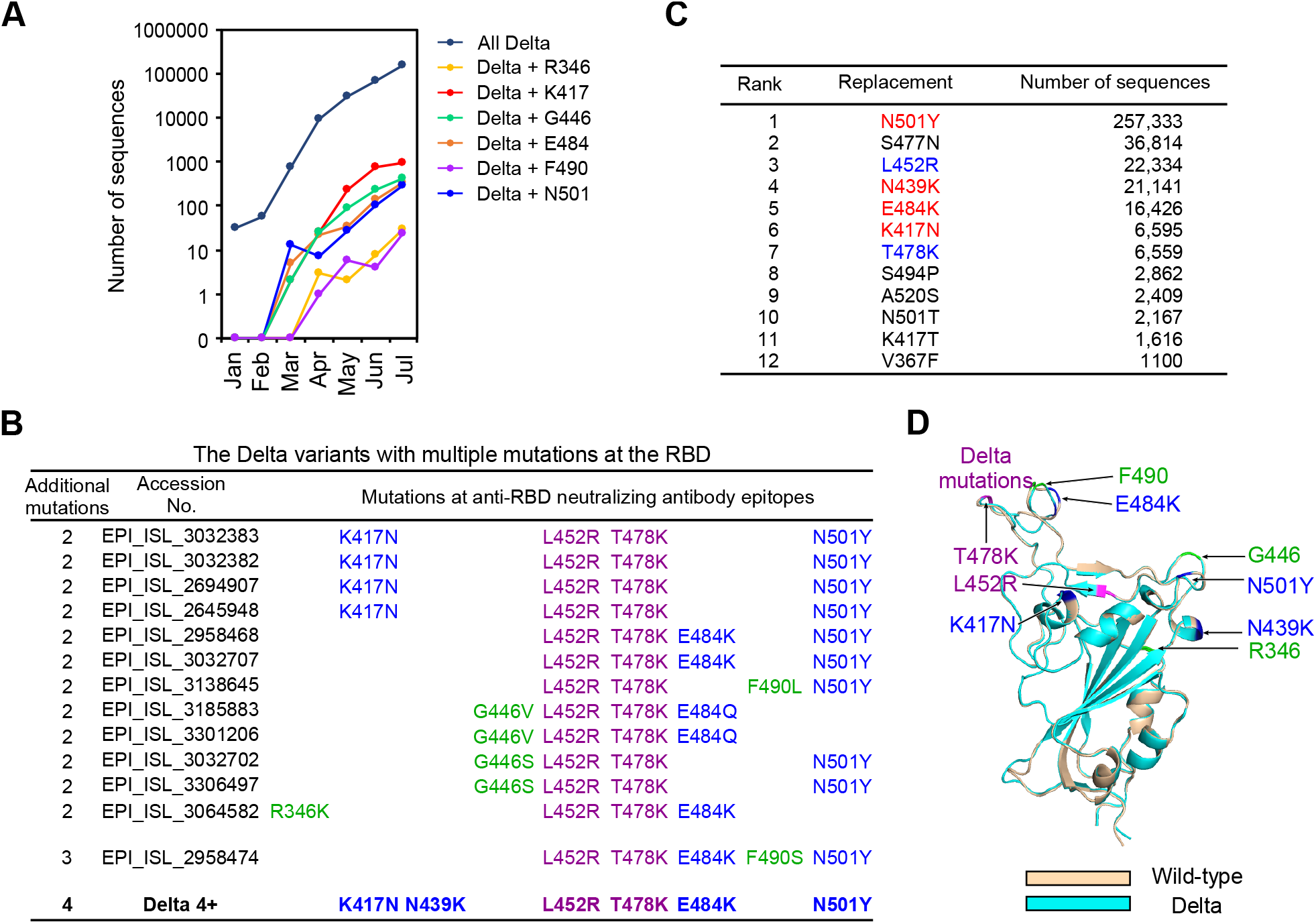
Possible mutations that may be acquired by the Delta variant. **(A)** Number of the Delta variants with additional mutations at the RBD registered in the GISAID database in each month from January, 2021 to July, 2021. The data registered at July are not enough and will be increased later. **(B)** The Delta variants with additional mutations at multiple epitopes of the anti-RBD neutralizing antibodies. L452R and T478K mutations are observed in all the Delta variants (purple). Anti-RBD neutralizing antibody epitopes introduced into the Delta 4+ (blue), and anti-RBD neutralizing antibody epitopes observed in the natural Delta variants but not introduced into the Delta 4+ (green) are shown with the respective GISAID accession number. **(C)** Number of the major RBD mutations acquired by all SARS-CoV-2 variants. L452R and T478 are mutations observed for the representative Delta variant (blue). N501Y, N439K, E484K and K417N were selected to generate the Delta 4+ variant (red). **(D)** Location of additional mutations introduced into the Delta RBD. Structures of the RBD of the wild-type (light brown) and the Delta variant (light blue) predicted by AlphaFold2 were superimposed. Mutations of the Delta variant (purple), anti-RBD neutralizing antibody epitopes to generate the Delta 4+ (blue), and anti-RBD neutralizing antibody epitopes observed in the natural Delta variants but not introduced into the Delta 4+ (shown in C; green) are indicated in the figure. See also Figure S4.

### Enhanced infectivity of the Delta 4+ pseudovirus by some BNT162b2-immune sera

We analyzed the binding of several anti-RBD neutralizing antibodies to the Delta spike with a single additional mutation or multiple mutations in the RBD (**Figure 6A**). Most anti-RBD antibodies recognized Delta spike with a single additional mutation, but not the Delta 4+ spike protein. The C135 anti-RBD neutralizing antibody, whose major epitopes are R346 and N440 (Greaney et al., 2021b; Weisblum et al., 2020), still recognized the Delta 4+ spike. We then generated pseudovirus bearing mutant spike proteins. The Delta pseudovirus with additional single RBD mutations was slightly more resistant to BNT162b2-immune sera (**Figure 6B**). The effects of the single additional mutations were slightly different depending on the individuals, although infection was completely blocked at the highest concentration of the serum. Next, we analyzed the Delta 4+ pseudovirus with four additional RBD mutations (**Figure 6C**). Surprisingly, most BNT162b2-immune sera enhanced infectivity of the Delta 4+ pseudovirus in a dose-dependent manner at relatively low concentrations of BNT162b2-immune sera, but showed weak neutralization only at the highest concentration of the sera (**Figure 6D and 6E**). Especially, PFZ7 greatly enhanced the infectivity at relatively low serum concentration. Some sera, such as PFZ13 and PFZ14, did not show neutralizing activity even at the highest concentration of the sera. The neutralizing titers of PFZ13 and PFZ14 against wild-type or Delta variant were apparently lower than others (**Figure 2A**). On the other hand, PFZ15 effectively neutralized the Delta 4+ pseudovirus, but the neutralizing titers of PFZ15 against the wild type and Delta variant were not particularly high compared to the others. Because most neutralizing antibodies against either NTD or RBD do not work for the Delta 4+ pseudovirus, while most enhancing antibodies remain functional for the Delta 4+ pseudovirus, the increased infectivity in the presence of BNT162b2-immune sera appears to be mediated by anti-NTD enhancing antibodies.

**Figure 6.**
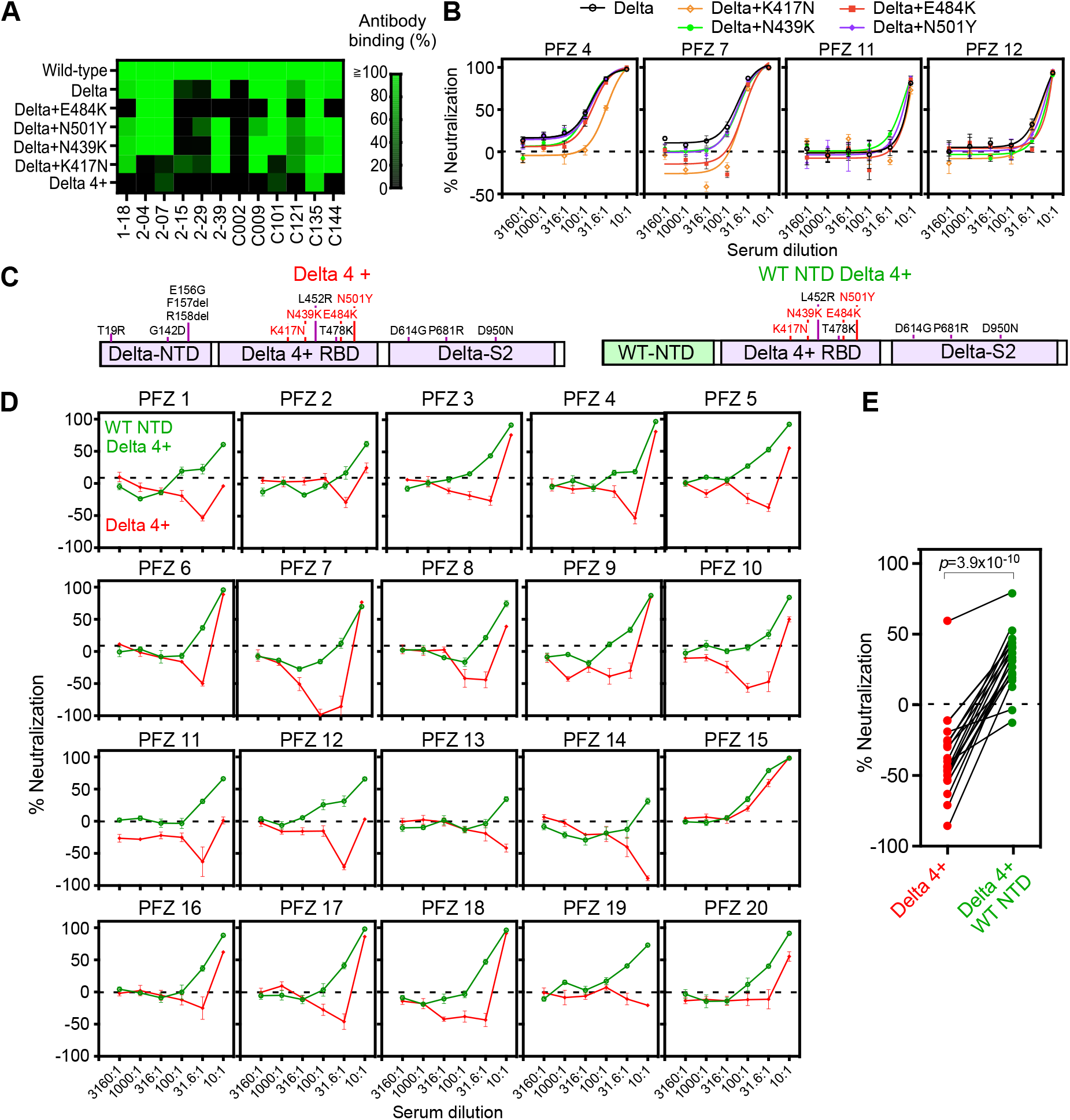
Enhanced infectivity of the Delta 4+ pseudovirus by the BNT162b2-immune sera. **(A)** Anti-RBD antibody binding to the Delta spike with additional mutations at the RBD. Anti-RBD mAb binding (1 μg/ml) to the mutant spike was compared to that of the wild-type spike. The Delta 4+ spike contains additional mutations of K417N, N439K, E484K and N501Y. **(B)** Neutralizing activity of BNT162b2-immune sera against the Delta pseudoviruses with a single additional mutation at the RBD as indicated in the figure. The data from quadruplicates are presented as mean ± SEM. **(C)** The construct of the Delta 4+ and Delta 4+ with wild-type (WT) NTD. Mutations in the original Delta variant (black) and the four mutations added to the Delta RBD (red) were shown. **(D)** Neutralizing activity of BNT162b2-immune sera against the pseudovirus with Delta 4+ spike (red) and Delta 4+ spike with wild-type NTD (green). **(E)** Neutralizing activity of 31.6 times diluted BNT162b2-immune sera. *p* value determined by paired t-test were indicated. Negative values for % neutralization indicates enhanced infectivity (B, D, E). The data from quadruplicates are presented as mean ± SEM. The representative data from three independent experiments are shown.

In order to analyze the contribution of Delta NTD to the enhanced infectivity, we generated pseudovirus bearing spike protein with wild-type NTD and Delta 4+ RBD (**Figure 6C)**. Although some BNT162b2-immune sera enhanced infectivity of the Delta 4+ pseudovirus, the Delta 4+ virus with wild-type NTD did not show enhanced infectivity by BNT162b2-immune sera (**Figure 6D and 6E**). These data suggested that mutations in the NTD of the Delta variant made the virus more susceptible than the wild-type to anti-NTD enhancing antibodies in BNT162b2-immune sera, and thus reduced the neutralizing effect of anti-RBD neutralizing antibodies.

### Sera from the Delta spike immunized mice do not show enhanced infectivity against Delta 4+ pseudovirus

Because wild-type spike was used for BNT162b2 mRNA vaccine, the enhanced infectivity of the Delta 4+ pseudovirus by some BNT162b2-immune sera appears to be caused by the decreased neutralizing antibody titer of anti-NTD and anti-RBD neutralizing antibodies against Delta 4+ pseudovirus. Therefore, neutralizing antibody titers against the Delta variants may be relatively high compared to enhancing antibodies when immunizing with the Delta spike, even though the enhancing antibody epitopes are conserved in the Delta spike protein. To test the effect of immunization by Delta spike, we immunized mice with B16F10 mouse melanoma cells transiently transfected with wild-type or Delta spike protein (**Figure 7A**). We used B16F10 cells because the immunogenicity of B16F10 melanoma cell line is quite low (Priem et al., 2020). In addition, the conformation of spike protein expressed on transfectants is likely to be similar to that of spike protein expressed by mRNA vaccines. All mice effectively produced antibodies against spike protein (**Figure S5**). The wild-type spike immunized sera neutralized wild-type pseudovirus well, whereas the neutralizing effect against the Delta pseudovirus decreased, similar to BNT162b2-immune sera (**Figure 7B and 7C**). In contrast, Delta spike immunized sera neutralized both wild-type and Delta pseudovirus well. Just one mouse produced antibodies that neutralize the Delta pseudovirus better than wild-type pseudovirus. When we analyzed the Delta-4+ pseudovirus, some sera from wild-type spike immunized mice showed enhanced infectivity in a dose dependent manner at relatively low concentrations of sera similar to some BNT162b2-immune sera (**Figure 7D and 7E**). Especially, #w1 mouse serum showed enhanced infectivity at any concentration, although the same serum neutralized the wild-type pseudovirus well. In contrast, the enhanced infectivity by immunized sera was not observed when the Delta spike was used for immunization. Sera from the Delta-spike immunized mice did not exhibit enhanced infectivity at any concentration of sera. These data suggest that vaccines containing the Delta, but not wild-type, spike might be required to control the Delta subvariant that may emerge in the future.

**Figure 7.**
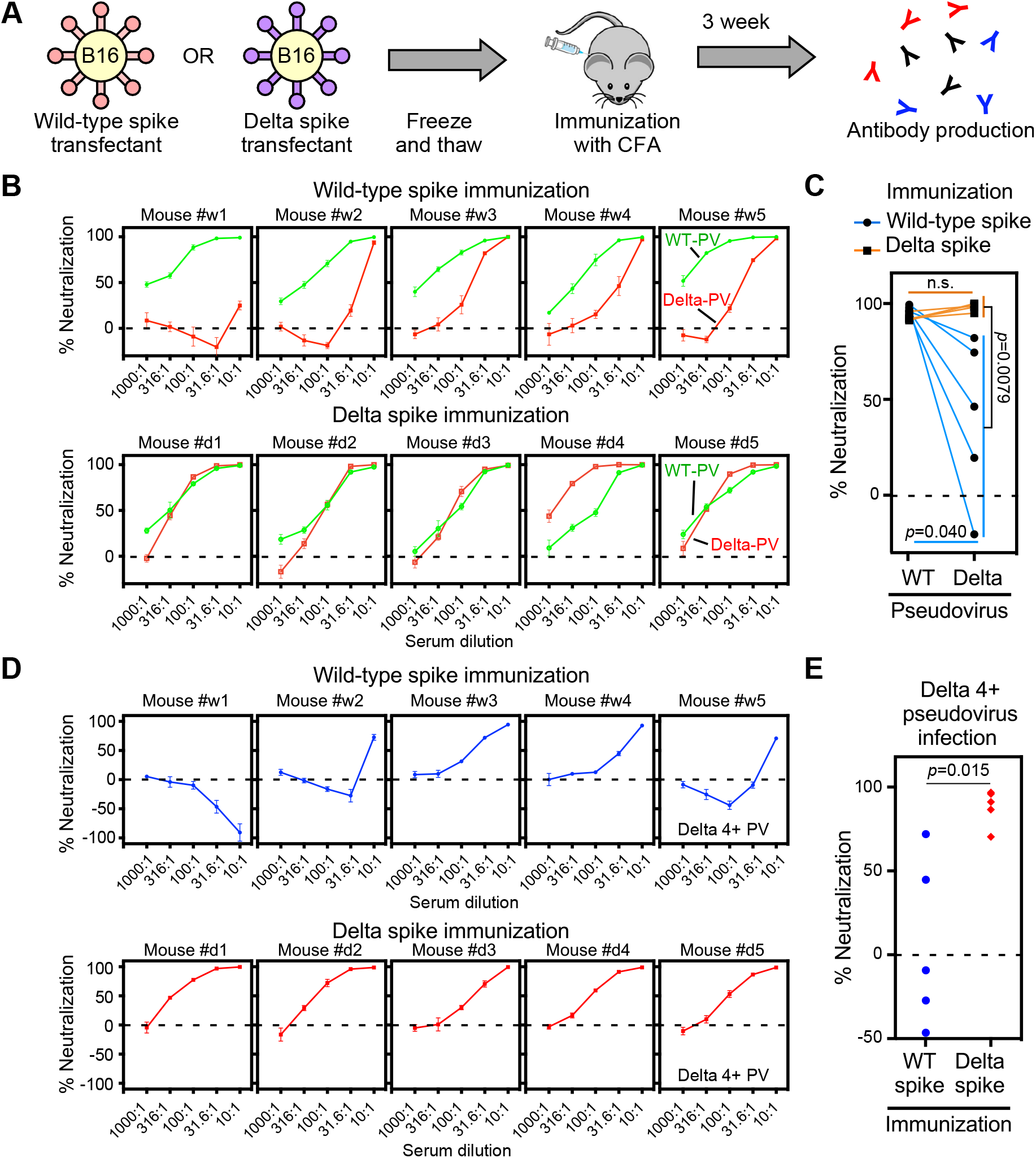
Sera from delta spike-immunized mice do not show enhanced infectivity. **(A)** Freeze and thawed wild-type and Delta spike-B16 transfectants were immunized to the mice with complete Freund’s adjuvant (CFA). **(B)** Neutralizing activity against the wild-type (green) or Delta (red) pseudovirus (PV) by sera from the wild-type spike (upper column) or Delta spike (lower column) spike-immunized mice. **(C)** Neutralizing activity against the wild-type and Delta pseudovirus by 31.6 times-diluted sera from wild-type (light blue line) or Delta (orange line) spike-immunized mice. **(D)** Neutralizing activity against the Delta 4+ pseudovirus by sera from the wild-type spike (upper column, blue) or Delta spike (lower column, red) immunized mice. **(E)** Neutralizing activity against the Delta 4+ pseudovirus by the 31.6 times-diluted sera from the wild-type spike (blue) or Delta spike (red) immunized mice. n.s.: not statistical significance, *p* value was determined by t-test. A negative values for % neutralization indicates enhanced infectivity. All data from quadruplicates are presented as mean ± SEM. See also Figure S1 and S5.

## Discussion

The Delta variant is highly contagious and breakthrough infection to fully vaccinated individuals is often observed (Lopez Bernal et al., 2021), suggesting that neutralizing antibodies in fully vaccinated individuals are not sufficient to protect against infection by the Delta variant. Anti-RBD antibodies are thought to play a major role in protection against SARS-CoV-2 infection. The Delta variant has L452R and T478K mutations in the RBD, and L452 has been shown to be an epitope for some neutralizing antibodies (McCallum et al., 2021; Wang et al., 2021b). However, most neutralizing antibodies bound to the Delta RBD and neutralized the infection. Therefore, mutations in the RBD alone may not explain the decreased neutralizing titers of the BNT162b2-immune sera against the Delta variant.

The Delta variant has multiple mutations in the NTD: T19R, G142D, E156G, F157del and R158del. All anti-NTD neutralizing antibodies failed to recognize the Delta spike, indicating that the Delta variant is completely resistant to anti-NTD neutralizing antibodies elicited by wild-type spike protein, which is the antigenic component of widely used mRNA vaccines. In contrast, most anti-NTD enhancing antibodies recognized Delta spike at the same level as wild-type spike, and some anti-NTD enhancing antibodies exhibited increased infectivity enhancement by Delta pseudovirus compared to wild-type pseudovirus. Consistent with this observation, the structures of enhancing anti-NTD antibody epitopes were well conserved with the wild type. Because enhancing antibodies reduced neutralizing activity of anti-RBD neutralizing antibodies (Li et al., 2021; Liu et al., 2021b), mutations in the NTD may play an important role in the resistance of the Delta variant to the BNT162b2-immune sera. Indeed, a Delta pseudovirus with wild-type NTD was more susceptible to neutralization by BNT162b2-immune sera than full Delta pseudovirus. The effect of the Delta NTD was more obvious for the Delta 4+ pseudovirus. These data indicated that mutations in the NTD are involved in the escape of SARS-CoV-2 from neutralizing antibodies. It is likely that the mutations in the NTD that abrogate neutralizing antibody binding while retaining enhancing antibody binding are beneficial to the virus. These mutations in the Delta variant may suggest adaptation to the presence of enhancing antibodies while maintaining evasion of anti-NTD and anti-RBD neutralizing antibodies in immunized or previously infected hosts.

Not only Delta, but also other VOCs such as Alpha (B.1.1.7), Beta (B.1.135), and Gamma (P.1) show more mutations in the NTD than in the RBD. Because the NTD is involved in the regulation of the conformation of the RBD but not in direct binding to the host receptor ACE2 (Liu et al., 2021b), it can tolerate many mutations. As with the Delta variant, most anti-NTD neutralizing antibodies have been reported not to bind to the Alpha and Beta variants (Voss et al., 2021; Wang et al., 2021a). Recently, L-SIGN has been reported to be an entry receptor for SARS-CoV-2 (Amraei et al., 2021; Kondo et al., 2021; Soh et al., 2020; Thepaut et al., 2021). L-SIGN specifically bound to NTD but not RBD of SARS-CoV-2 spike protein and mediated SARS-CoV-2 infection of non-ACE2 expressing cells by inducing membrane fusion (Soh et al., 2020). Furthermore, anti-NTD neutralizing antibodies efficiently blocked SARS-CoV-2 infection of L-SIGN-expressing cells compared to that of ACE2-expressing cells. Considering the fact that most VOCs have completely escaped from anti-NTD neutralizing antibodies regardless of the fact that the neutralizing efficiency is quite low compared to anti-RBD neutralizing antibodies *in vitro*, SARS-CoV-2 infection mediated by the NTD through L-SIGN or other unknown receptors may play a more important role *in vivo* than *in vitro*. Further analyses of function of NTD as well as anti-NTD neutralizing antibodies are required to elucidate the pathogenicity of SARS-CoV-2.

The enhancing antibodies bind to a specific site on the NTD, inducing the open form of the RBD, which increases the affinity of spike protein to ACE2 (Liu et al., 2021b). Recently, it has been reported that the enhancing antibodies do not increase the infectivity *in vivo* (Li et al., 2021). However, only one human IgG1 monoclonal enhancing antibody, among 11 known enhancing antibodies, has been tested *in vivo*. The affinities and epitopes of enhancing antibodies to the NTD, as well as the IgG subclass of enhancing antibodies, may affect their *in vivo* function. Recently, it has been reported that binding of neutralizing antibodies to Fc receptors is required for their neutralizing activity *in vivo* (Schafer et al., 2021; Suryadevara et al., 2021; Winkler et al., 2021). Indeed, IgG1, which is the most frequently used antibody subclass in *in vivo* studies, has the strongest affinity for Fc receptors and shows strong effector function; whereas, IgG2 and IgG4 weakly bind to Fc receptors (Nimmerjahn and Ravetch, 2008). Therefore, it is likely that the *in vivo* function of anti-NTD enhancing antibodies will vary depending on the antibody subclass, the specific variable region sequence, or both. Given the fact that the Delta variant maintained enhancing antibody epitopes and is more sensitive to enhancing antibodies, it is likely that the enhancing antibodies are involved in augmentation of the SARS-CoV-2 infectivity *in vivo*.

Several BNT162b2 immune sera showed neutralizing activity against the Delta 4+ pseudovirus at a 1:10 dilution, but conversely increased infectivity at 1:30 dilution. In general, the activity of neutralizing antibodies does not change so drastically with a three-fold difference in concentration. Therefore, the effect of the BNT162b2 immune sera against the Delta 4+ pseudovirus cannot be explained simply by the concentration of neutralizing antibodies. The BNT162b2 immune sera did not show enhanced infectivity against the Delta 4+ pseudovirus with wild-type NTD at any serum concentration. Since the effect of anti-NTD infectivity-enhancing monoclonal antibodies is affected by the concentration of anti-RBD neutralizing antibodies (Li et al., 2021; Liu et al., 2021b), the effect of infectivity-enhancing antibodies in BNT162b2 immune sera is likely to be more pronounced when the concentration of anti-RBD neutralizing antibodies falls below a certain threshold. Indeed, the BNT162b2 immune sera with low neutralizing titers against the Delta pseudovirus showed enhancement against the Delta 4+ pseudovirus even at high serum concentration. Although the neutralizing antibody titer is the highest three weeks after the second immunization, it gradually decreases (Doria-Rose et al., 2021; Widge et al., 2021). As in the case of diluted sera, it is possible that the effect of infectivity-enhancing antibodies may become more evident some time after immunization, even if the neutralizing and enhancing antibody titers decrease equally. In addition, neutralizing antibody titers induced by adenovirus vaccines and inactivated vaccines are lower than those induced by mRNA vaccines (Lim et al., 2021; Shrotri et al., 2021). Therefore, there is a possibility that the enhancing effect might be more pronounced against the Delta 4+ pseudovirus with immune sera of adenovirus vaccines or inactivated vaccines, similar to BNT162b2 immune sera with low neutralizing titers. On the other hand, some BNT162b2 immune sera did not enhance infection of Delta 4+ pseudovirus at any serum concentration and neutralized well. Similarly, despite the use of inbred mice, the effect of sera on the infectivity of Delta 4+ pseudovirus varied greatly among individual mice immunized with the wild-type spike. The sera of some mice showed enhancement of the Delta 4+ pseudovirus infection, while others showed neutralization at any serum concentration. The delicate balance of antibody titer, affinity, or epitope between neutralizing and enhancing antibodies might affect the effect of sera on the infectivity. It is important to further analyze the characteristics of neutralizing and enhancing antibodies produced after immunization.

SARS-CoV-2 has acquired a number of mutations to date, which have arisen within infected individuals. Therefore, new variants are likely to emerge more frequently in situations where many people are infected. Because the Delta variant is spreading so explosively, it has already acquired numerous additional mutations in the spike protein coding region, suggesting that the Delta variant will continue to acquire further mutations. Some mutations observed in the RBD of the Delta variant have been reported to be epitopes for anti-RBD neutralizing antibodies (Greaney et al., 2021a; Greaney et al., 2021b; Wang et al., 2021b). Newly emerged variants that adapt to the environment of their host’s immune system will be selected and expand. The Delta variant with 4 additional mutations in the RBD were not neutralized by most BNT162b2-immune sera because of unique mutations in the NTD. More importantly, infectivity of the Delta 4+ was enhanced by some BNT162b2-immune sera. Furthermore, of the four additional mutations, a Delta variant with three mutations has already been registered in the GISAID database; it is likely that a Delta variant that has acquired five mutations in the RBD in total will acquire additional mutations in the near future. Although we have selected K417N, N439K, E484K, and N501Y as additional mutations for the Delta variant, other combinations of anti-RBD neutralizing epitopes can be expected to have similar or stronger effects than the Delta 4+ variant. Indeed, the Delta 4+ still possess R346, one of major epitope residues for anti-RBD neutralizing antibodies such as C135. Given the current high mutation rate of SARS-CoV-2, predicting emerging spike mutations is very important to develop effective vaccines against emerging SARS-CoV-2 variants. Immunization by dangerous spike protein variants that are likely to emerge in the future may be effective in preventing the emergence of such variants.

A third round of booster immunization with the SARS-CoV-2 vaccine is currently under consideration. Our data suggest that repeated immunization with the wild-type spike may not be effective in controlling the newly emerging Delta variants. We demonstrated that immunization by Delta spike induces antibodies that neutralize not only the Delta variant but also wild-type and the Delta 4+ variant without enhancing the infectivity. Although mRNA vaccination may yield different results from our animal model, development of mRNA vaccine expressing the Delta spike might be effective for controlling the emerging Delta variant. However, epitopes of the enhancing antibodies, not neutralizing antibodies, are well conserved in most SARS-CoV-2 variants, including the Delta variant. Therefore, additional immunization of the spike protein derived from SARS-CoV-2 variants may boost enhancing antibodies more than the neutralizing antibodies in individuals who were previously infected with wild-type SARS-CoV-2 or immunized with vaccines composed of wild-type spike protein. Immunization using the RBD alone, which will not induce anti-NTD enhancing antibodies, could be a strategy for a vaccination. However, anti-NTD neutralizing antibodies that protect against SARS-CoV-2 infection similar to anti-RBD-neutralizing antibodies are not induced by immunization by RBD alone (Chi et al., 2020; Li et al., 2021; Liu et al., 2020; Suryadevara et al., 2021; Voss et al., 2021). Whole spike protein containing RBD mutations observed in major variants but lacking the enhancing antibody epitopes may need to be considered as a booster vaccine.

## Acknowledgments

We would like to thank Yumi Inaba for her administrative assistance. We would like to thank technical assistance of the members of the Department of Dermatology, Osaka University, Osaka, Japan. This work was supported by JSPS KAKENHI under Grant Numbers JP18H05279 and JP19H03478, MEXT KAKENHI under Grant Number JP19H04808, Japan Agency for Medical Research and Development (AMED) under Grant Numbers, JP20fk0108542 (HA), JP20nf0101623 (HN, HA), 20fk0108403h (HA), JP20am0101108 (DMS), JP21am0101072 (TK, AN, BINDS support number 2630) and Japan Science and Technology Agency (JST) (Moonshot R&D) JPMJMS2025 (YM), and Panasonic Corporation (HA, YM).

## Author contributions

Y.L., Y.M., D.M.S., T.K., M.O., M.F., H.A. designed the experiments. Y.L., J.K., M.H., A.T, S.M., A.A., K.A., C.O., H.J., K.K., W.N., performed the experiments. N.A., A.K., H.N., Y.Y., M.F. collected vaccine sera. J.K., S.L., D.M.S, T.K., H.A. constructed a model of NTD spike. Y.L., N.A., J.K, M.K., D.M.S, H.A. wrote the manuscript. All authors read, edited, and approved the manuscript.

## Methods

### Data and code availability

Cryo-EM density maps for the SARS-CoV-2 Delta spike protein were deposited at the EMDB under accession code EMD-31731. A molecular model of the SARS-CoV-2 Delta spike protein fitted to Cryo-EM data were deposited to PDB under accession code 7V5W. The data that support the findings of this study are available from the Lead Contact on request.

### Cell lines

HEK293T cells (RIKEN Cell Bank) and B16F10 melanoma cells (National Institute of Biomedical Innovation) were cultured in DMEM (Nacalai, Japan) supplemented with 10% FBS (Biological Industries, USA), penicillin (100 U/mL), and streptomycin (100 μg/mL) (Nacalai, Japan) and cultured at 37°C in 5% CO_2_. The Expi293 cells (Thermo) were cultured with the Expi293 medium. The cells were routinely checked for mycoplasma contamination. ACE2-stably transfected HEK293 cells (HEK293T-ACE2-transfectants) were reported previously (Liu et al., 2021b).

### Human samples

The collection and use of BNT162b2-immune sera were approved by Osaka University Hospital (20522-3). Written informed consent was obtained from the participants according to the relevant guidelines of the institutional review board. All sera were collected from 26-65 years old healthy individuals three weeks after immuniztion with two cycles of 30 μg of BNT162b2 mRNA vaccine.

### Plasmid construction

The SARS-CoV-2 spike gene (NC_045512.2) was prepared by gene synthesis (IDT). The sequences encoding the spike protein lacking the C-terminal 19 amino acids (amino acids 1–1254) were cloned into the pME18S expression vector. NTD (amino acids 14–333) and RBD (amino acids 335–587) were separately cloned into a pME18S expression vector containing a SLAM signal sequence and a PILRα transmembrane domain (Saito et al., 2017). A series of mutants and the Delta variants (T19R, G142D, E156G, del_157, del_158, L452R, T478K, D614G, P681R, D950N) were prepared from wild-type SARS-CoV-2 spike using the QuickChange Lighting Multi Site-directed Mutagenesis kit (Agilent). Additional RBD mutations were introduced into the Delta spike also using the QuickChange Lighting Multi Site-directed Mutagenesis kit (Agilent). The primers for mutagenesis were designed on Agilent’s website (https://www.agilent.com/store/primerDesignProgram.jsp). For Cryo-EM analysis, the sequence encoding the spike protein’s extracellular domain with a foldon and His-tag at the C-terminus (Cai et al., 2020) was cloned into a pcDNA3.4 expression vector containing the SLAM signal sequence. Also, mutations D614G, R686G R687S R689G, K986P, and V987P were introduced using a Quick change multi-mutagenesis kit (Agilent) for stabilization of recombinant spike protein (Yurkovetskiy et al., 2020). The DNA sequences of these constructs were confirmed by sequencing (ABI3130xl).

### Transfection

A pME18S expression plasmid containing the full-length or subunit spike protein was transiently transfected into HEK293T cells using PEI max (Polysciences); the pMx-GFP expression plasmid was used as the marker of transfected cells.

### Anti-spike monoclonal antibodies from COVID-19 patients

The variable regions of anti-SARS-CoV-2 spike antibodies from COVID-19 patients were synthesized according to the published sequence (IDT) (Brouwer et al., 2020; Chi et al., 2020; Li et al., 2021; Robbiani et al., 2020; Suryadevara et al., 2021; Zost et al., 2020). Variable region sequences of some antibodies were obtained from the CoV-AbDab database (http://opig.stats.ox.ac.uk/webapps/covabdab/). The cDNA of the variable regions of the heavy chain and light chain were cloned into a pCAGGS vector containing sequences that encode the human IgG1 or kappa constant region. The pCAGGS vectors containing sequences encoding the immunoglobulin heavy chain and light chain were co-transfected into Expi293 (Thermo) cells, and the cell culture supernatants were collected according to the manufacturer’s protocols. Recombinant IgG was purified from the culture supernatants using protein A Sepharose (GE healthcare). The concentration of purified IgG was measured at OD280.

### Antibodies and recombinant proteins

Allophycocyanin (APC)-conjugated donkey anti-mouse IgG Fc fragment antibody and APC-conjugated anti-human IgG Fc fragment specific antibody (Jackson ImmunoResearch, USA) were used. The pcDNA3.4 expression vector containing the sequence that encodes the His-tagged extracellular domain of the spike protein was transfected into Expi293 cells and the His-tagged spike protein produced in the culture supernatants was then purified with a Talon resin (Clontech).

### Immunization of mice

B16F10 cells were transfected with WT spike protein or Delta spike protein by PEI as described above. 48 hours later, B16F10 cells were washed twice with PBS,and then the cells were collected and frozen and thawed. Balb/c female mice (7-weeks-old females) were purchased from SLC. Two groups of five mice (n = 5) were subcutaneously immunized with 1×10^7^ B16F10 transfectants in the presence of complete Freund’s adjuvant (CFA). Serum samples were collected three weeks after the immunization.

### Flow cytometric analysis of antibodies

Plasmids expressing the full-length SARS-CoV-2 spike protein, Flag-NTD-PILR-TM and Flag-RBD-PILR-TM were co-transfected with the GFP vector into HEK293T cells. The transfectants were incubated with the mAbs, followed by APC-conjugated anti-human IgG Ab. The antibodies bound to the stained cells were then analyzed using a flow cytometer (Attune^™^, Thermo; FACSCelesta BD bioscience). Antibodies binding to the GFP-positive cells were shown in the figures using FlowJo software (BD bioscience).

### SARS-CoV-2 spike-pseudotyped virus infection assay

The HEK293T cells were transiently transfected with expression plasmids for the SARS-CoV-2 spike protein lacking the C-terminal 19 amino acids (Hu et al., 2020; Johnson et al., 2020). At 24 hours post-transfection, VSV-G-deficient VSV carrying a Luciferase gene complemented in *trans* with the VSV-G protein was added for incubation for 2 hours. The cells were then carefully washed with DMEM media without FBS and incubated with DMEM with FBS at 37°C in 5% CO_2_ for 48 hours. The supernatant containing the pseudotyped SARS-CoV-2 virions was harvested and aliquoted before storage at −80°C. To determine the virus titers of the pseudovirus, 1× 10^4^ HEK293T-ACE2-transfectants were mixed with the pseudovirus for 20 hours at 37 °C in 5% CO_2_ in a 384-well plate (Greiner, Germany). Luciferase activity was measured using a ONE-Glo^™^ luciferase assay (Promega, USA) according to the manufacturer’s instructions. The signals were measured by a luminescence plate reader (TriStar LB94, Berthold Technologies, Germany) **(Figure S1)**. For the neutralization assay, 5 μl pseudovirus was mixed with equal volume of sera or monoclonal antibodies at the concentrations indicated in the figure. The mixture was added to 20 μl of 1× 10^4^ HEK293T-ACE2-transfectants. To calculate % neutralization, the relative luminescence units of the virus control wells (pseudovirus only) were subtracted from those of the sample wells, and the subtaracted values were divided by those of the virus control wells. The PRNT50 neutralization titers for vaccinated sera were determined using 3-parameter nonlinear regression curve (GraphPad Prism). If the PRNT50 titer was less than 1:10, it was defined as 0.

### Structure prediction by AlphaFold2

The NTD and RBD structures of the wild type and Delta variant were predicted by AlphaFold2 (Jumper et al., 2021). The structure of the NTD was predicted in CASP14 mode without template. The structure of the RBD was predicted in CASP14 mode, using the template of 2020-05-14. The highest ranked prediction results were used.

### Cryo-EM data collection

A 2.5 μl protein solution of the spike protein (2.2 mg/ml) was applied onto the cryo-grid and frozen in liquid ethane using a Vitrobot IV (Thermo Fisher Scientific, USA, 4°C and 100% humidity). Quantifoil Au R0.6/1.0 holey carbon grids were used for the grid preparation. Data collection of the sample was carried out on a Titan Krios (Thermo Fisher Scientific, USA) equipped with a thermal field emission electron gun operated at 300 kV, an energy filter with a 20 eV slit width and a bioquantum K3 direct electron detection camera (Gatan, USA) (Figure S4). For automated data acquisition, SerialEM software was used to collect cryo-EM image data. Movie frames were recorded using the K3 camera at a calibrated magnification of × 81,000 corresponding to a pixel size of 0.88 Å with a setting defocus range from −0.8 to −2.0 μm. The data were collected with a total exposure of 3 s fractionated into 62 frames, with a total dose of ~ 60 electrons Å^2^ in counting mode. A total number of movies were collected; 15,000 for the spike protein.

### Image processing and 3D reconstruction

All of image processes were carried out on cryoSPARC software (Punjani et al., 2017). After motion correction of movies and CTF parameter estimation, the particles were automatically picked using Topaz software (Bepler et al., 2019). The detailed information is summarized in Table S1. The picked particles were extracted into a box of 360 × 360 pixels. After particle extraction, the particles were applied to two rounds of heterogenous refinement with C1 symmetry. The selected particles (735,623 particles) were applied to two rounds of *ab-initio* reconstruction into three classes with C1 symmetry. In the first and second rounds of ab-initio reconstruction, the class similarity parameter, 0.1 and 0.8, was used, respectively. After that, the selected 147,497 particles were further used as non-uniform refinement with optimizing per-particle defocus. As the result, the density map for the spike protein was obtained at 3.16 Å resolution. Local resolution of the obtained maps were estimated by Local resolution estimation job on cryoSPARC.

### Model building and refinement

To generate the atomic model for the spike protein, the structure of NTD of Delta variant was predicted using AlphaFold2 (Jumper et al., 2021). For other domains, the model from previous study (PDBID; 7JJI) was used. These structures were fitted into the density map as rigid body using UCSF chimera (Pettersen et al., 2004). The initial model was extensively manually corrected residue by residue in COOT (Emsley et al., 2010) in terms of especially side-chain conformations. The corrected model was refined by the phenix.real_space_refine program (Liebschner et al., 2019) with secondary structure and Ramachandran restraints, then the resulting model was manually checked by COOT. This iterative process was performed for several rounds to correct remaining errors until the model was in good agreement with geometry, as reflected by the MolProbity score of 2.07 (Williams et al., 2018). For model validation against over-fitting, the built models were used for calculation of FSC curves against the final density map used for model building by phenix.refine program. The statistics of the obtained maps and the atomic model were summarized in Supplemental Table S1.

### Data and statistical analysis

FlowJo version 10.7 (BD Biosciences, USA) was used to analyze the flow cytometry data, and Graphpad Prism version 7.0e was used for graph generation and statistical analysis.

---

**Figure S1.**
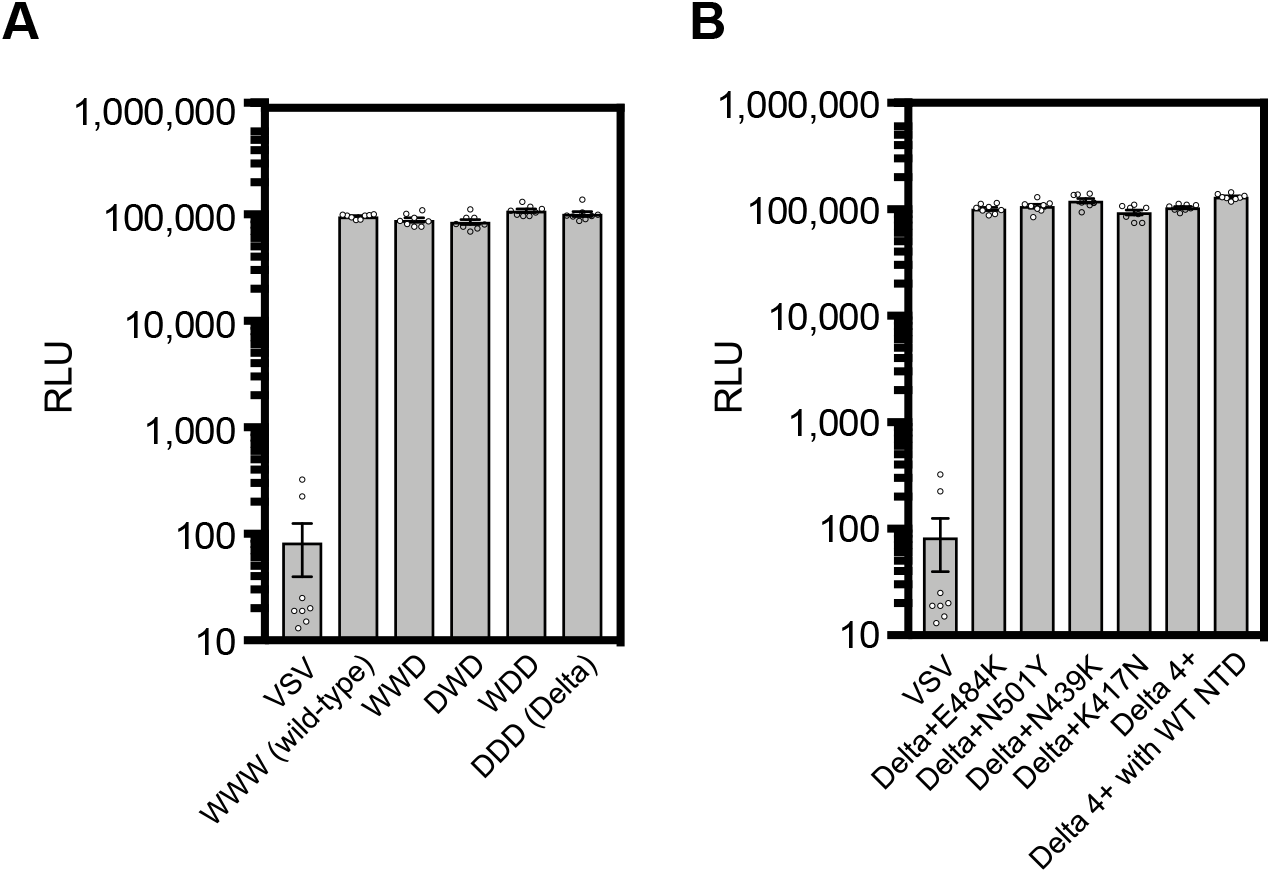
Viral titers of pseudotyped viruses, related to Figure 1, 2, 3, 6 and 7. The viral titer for each psudoviruses was measured by infection of ACE2-transfected HEK293T cells as described in Methods.

**Figure S2.**
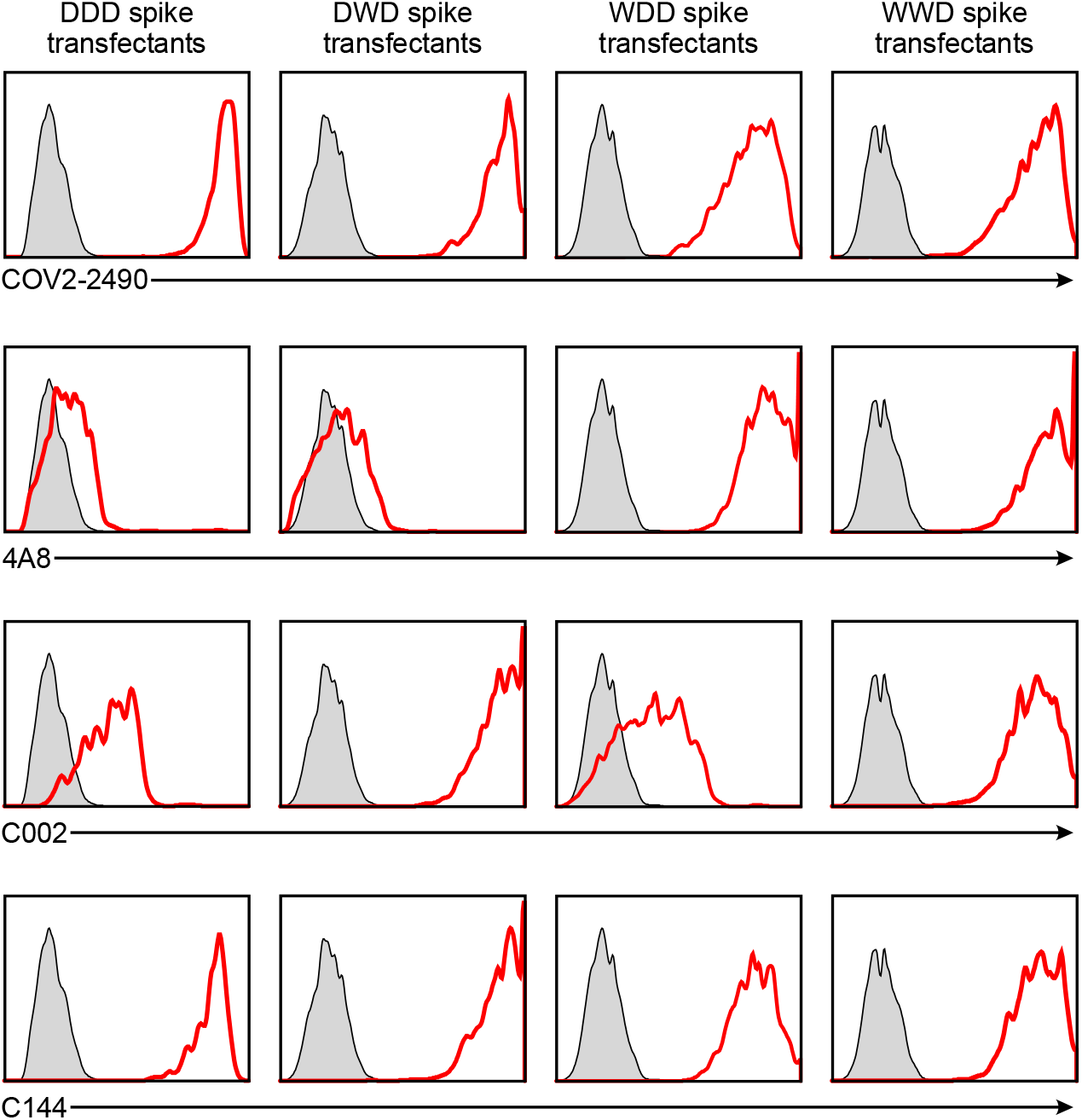
Anti-spike monoclonal antibody binding to the chimeric spike proteins, related to Figure 3. Chimeric spike proteins DDD, DWD, WDD and WWD were transfected with GFP to HEK293T cells and the transfectants were stained with 1 μg/ml COV2-2490, 4A8, C002, and C144 antibodies. Antibody bound to the GFP positive cells are shown (red histogram). Control staining: shaded histogram.

**Figure S3.**
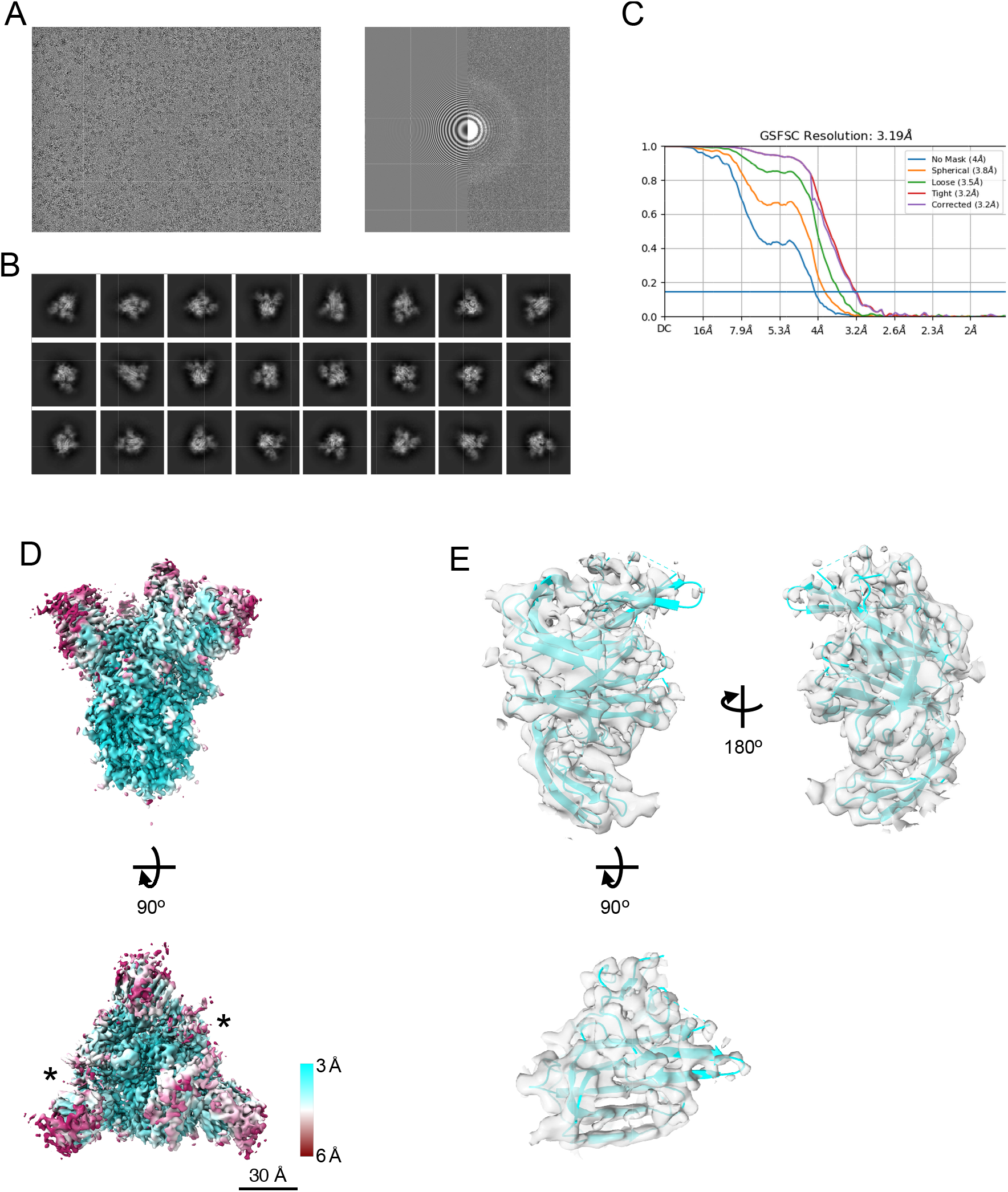
Cryo-EM density map of spike of SARS-CoV-2 Delta variant, related to Figure 4. **(A)** A representative micrographs (left), CTF estimation of a micrograph on left panel (right). **(B)** Typical 2D class averages. **(C)** The GS-FSC curves for the obtained map from cryoSPARC software are shown. Blue flat line indicates FSC=0.143 criteria. **(D)** The density map of spike protein from Delta strain (EMDBID: 31731). The map is colored with local resolution. Asterisks indicate the up form of RBDs. Scale bars are 30 Å. **(E)** The structure of NTD from spike protein of Delta variant. The density map and the model are shown as semi-transparent surface and cartoon, respectively (PDBID: 7V5W).

**Figure S4.**
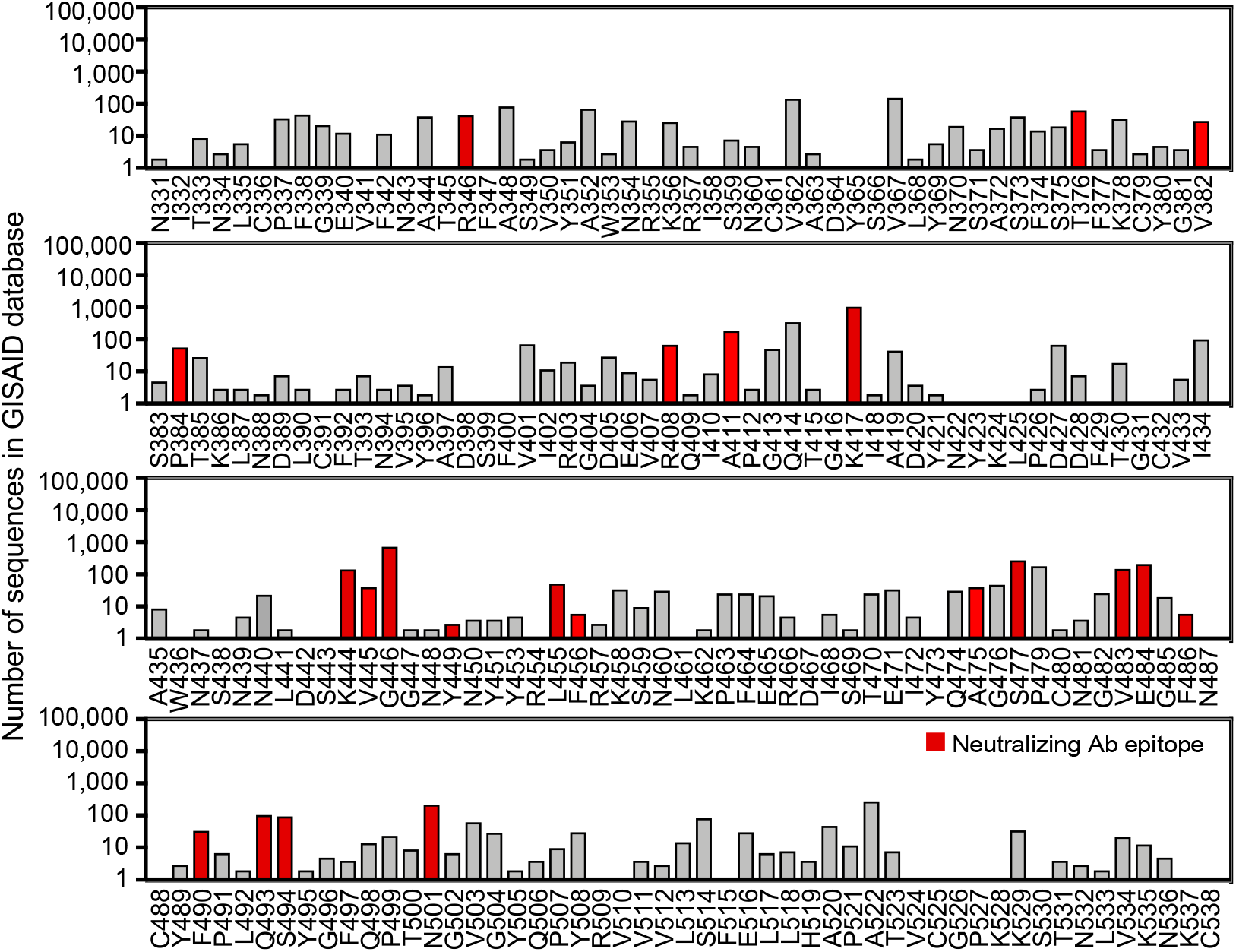
RBD mutations acquired by the Delta variant, related to Figure 5. Number of RBD mutations acquired by the Delta variant. The numbers of mutations at each residue registered in the GISAID database are shown. L452 and T478 mutations included in all the Delta variant were excluded. The red bars indicate the known epitopes for anti-RBD neutralizing antibodies.

**Figure S5.**
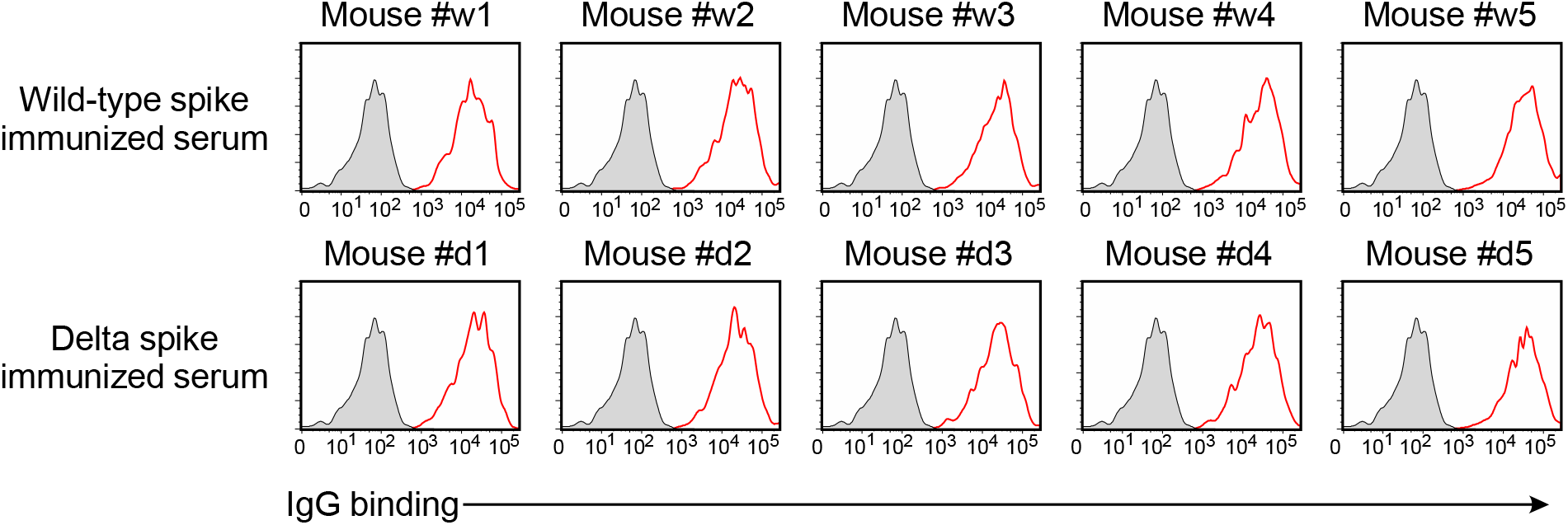
Anti-spike antibodies of the wild-type and delta spike-immunized mice, related to Figure 7. IgG antibody binding of the 100 times diluted spike-immunized mouse sera to the wild-type spike transfectants were analyzed by flow cytometer. Red: IgG binding. Gray: Control staining.

**Table S1.**
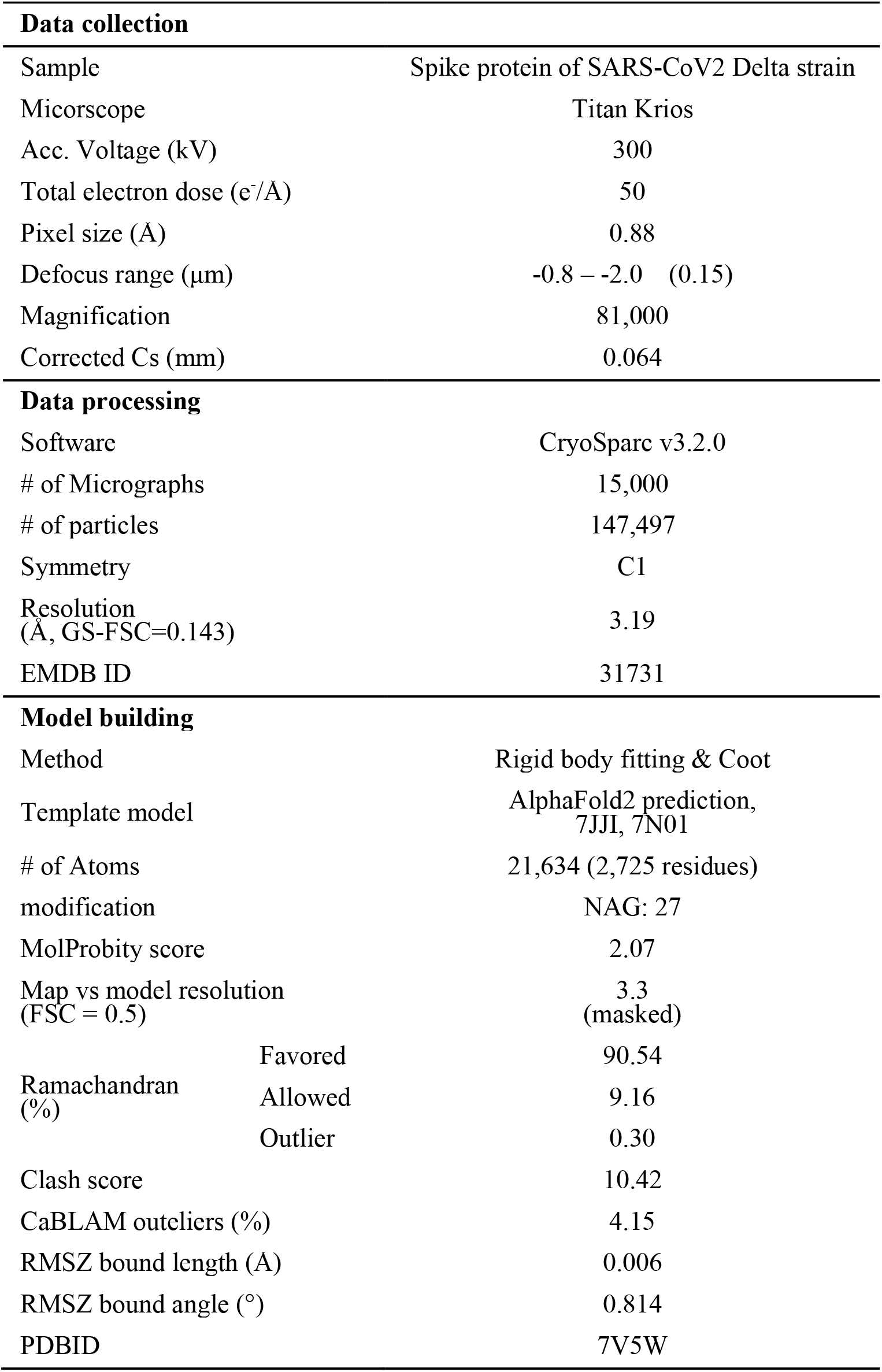
Cryo-EM data collection and processing statistics, related to Figure 4.

## References

Amraei, R., Yin, W., Napoleon, M.A., Suder, E.L., Berrigan, J., Zhao, Q., Olejnik, J., Chandler, K.B., Xia, C., Feldman, J., et al. (2021). CD209L/L-SIGN and CD209/DC-SIGN act as receptors for SARS-CoV-2. bioRxiv 10.1101/2020.06.22.165803.

Bepler, T., Morin, A., Rapp, M., Brasch, J., Shapiro, L., Noble, A.J., and Berger, B. (2019). Positive-unlabeled convolutional neural networks for particle picking in cryo-electron micrographs. Nat Methods 16, 1153–1160.

Brouwer, P.J.M., Caniels, T.G., van der Straten, K., Snitselaar, J.L., Aldon, Y., Bangaru, S., Torres, J.L., Okba, N.M.A., Claireaux, M., Kerster, G., et al. (2020). Potent neutralizing antibodies from COVID-19 patients define multiple targets of vulnerability. Science 369, 643–650.

Cai, Y., Zhang, J., Xiao, T., Peng, H., Sterling, S.M., Walsh, R.M., Jr., Rawson, S., Rìts-Volloch, S., and Chen, B. (2020). Distinct conformational states of SARS-CoV-2 spike protein. Science 369, 1586–1592.

Callaway, E. (2021). Delta coronavirus variant: scientists brace for impact. Nature 595, 17–18.

Cele, S., Gazy, I., Jackson, L., Hwa, S.H., Tegally, H., Lustig, G., Giandhari, J., Pillay, S., Wilkinson, E., Naidoo, Y., et al. (2021). Escape of SARS-CoV-2 501Y.V2 from neutralization by convalescent plasma. Nature 593, 142–146.

Chi, X., Yan, R., Zhang, J., Zhang, G., Zhang, Y., Hao, M., Zhang, Z., Fan, P., Dong, Y., Yang, Y., et al. (2020). A neutralizing human antibody binds to the N-terminal domain of the Spike protein of SARS-CoV-2. Science 369, 650–655.

Collier, D.A., De Marco, A., Ferreira, I., Meng, B., Datir, R.P., Walls, A.C., Kemp, S.A., Bassi, J., Pinto, D., Silacci-Fregni, C., et al. (2021). Sensitivity of SARS-CoV-2 B.1.1.7 to mRNA vaccine-elicited antibodies. Nature 593, 136–141.

Davies, N.G., Abbott, S., Barnard, R.C., Jarvis, C.I., Kucharski, A.J., Munday, J.D., Pearson, C.A.B., Russell, T.W., Tully, D.C., Washburne, A.D., et al. (2021). Estimated transmissibility and impact of SARS-CoV-2 lineage B.1.1.7 in England. Science 372.

Doria-Rose, N., Suthar, M.S., Makowski, M., O’Connell, S., McDermott, A.B., Flach, B., Ledgerwood, J.E., Mascola, J.R., Graham, B.S., Lin, B.C., et al. (2021). Antibody Persistence through 6 Months after the Second Dose of mRNA-1273 Vaccine for Covid-19. N Engl J Med 384, 2259–2261.

Emsley, P., Lohkamp, B., Scott, W.G., and Cowtan, K. (2010). Features and development of Coot. Acta Crystallogr D Biol Crystallogr 66, 486–501.

Greaney, A.J., Loes, A.N., Gentles, L.E., Crawford, K.H.D., Starr, T.N., Malone, K.D., Chu, H.Y., and Bloom, J.D. (2021a). Antibodies elicited by mRNA-1273 vaccination bind more broadly to the receptor binding domain than do those from SARS-CoV-2 infection. Sci Transl Med 13, eabi9915.

Greaney, A.J., Starr, T.N., Barnes, C.O., Weisblum, Y., Schmidt, F., Caskey, M., Gaebler, C., Cho, A., Agudelo, M., Finkin, S., et al. (2021b). Mapping mutations to the SARS-CoV-2 RBD that escape binding by different classes of antibodies. Nat Commun 12, 4196.

Greaney, A.J., Starr, T.N., Gilchuk, P., Zost, S.J., Binshtein, E., Loes, A.N., Hilton, S.K., Huddleston, J., Eguia, R., Crawford, K.H.D., et al. (2021c). Complete Mapping of Mutations to the SARS-CoV-2 Spike Receptor-Binding Domain that Escape Antibody Recognition. Cell Host Microbe 29, 44–57 e49.

Gupta, N., Kaur, H., Yadav, P., Mukhopadhyay, L., Sahay, R.R., Kumar, A., Nyayanit, D.A., Shete, A.M., Patil, S., Majumdar, T., et al. (2021). Clinical characterization and Genomic analysis of COVID-19 breakthrough infections during second wave in different states of India. medRxiv 10.1101/2021.07.13.21260273.

Hoffmann, M., Kleine-Weber, H., Schroeder, S., Kruger, N., Herrler, T., Erichsen, S., Schiergens, T.S., Herrler, G., Wu, N.H., Nitsche, A., et al. (2020). SARS-CoV-2 Cell Entry Depends on ACE2 and TMPRSS2 and Is Blocked by a Clinically Proven Protease Inhibitor. Cell 181, 271–280 e278.

Hu, J., Gao, Q., He, C., Huang, A., Tang, N., and Wang, K. (2020). Development of cell-based pseudovirus entry assay to identify potential viral entry inhibitors and neutralizing antibodies against SARS-CoV-2. Genes Dis 7, 551–557.

Jackson, L.A., Anderson, E.J., Rouphael, N.G., Roberts, P.C., Makhene, M., Coler, R.N., McCullough, M.P., Chappell, J.D., Denison, M.R., Stevens, L.J., et al. (2020). An mRNA Vaccine against SARS-CoV-2 - Preliminary Report. N Engl J Med 383, 1920–1931.

Johnson, M.C., Lyddon, T.D., Suarez, R., Salcedo, B., LePique, M., Graham, M., Ricana, C., Robinson, C., and Ritter, D.G. (2020). Optimized Pseudotyping Conditions for the SARS-COV-2 Spike Glycoprotein. J Virol 94, e01062–01020.

Jumper, J., Evans, R., Pritzel, A., Green, T., Figurnov, M., Ronneberger, O., Tunyasuvunakool, K., Bates, R., Zidek, A., Potapenko, A., et al. (2021). Highly accurate protein structure prediction with AlphaFold. Nature 10.1038/s41586-021-03819-2.

Kondo, Y., Larabee, J.L., Gao, L., Shi, H., Shao, B., Hoover, C.M., McDaniel, J.M., Ho, Y.C., Silasi-Mansat, R., Archer-Hartmann, S.A., et al. (2021). L-SIGN is a receptor on liver sinusoidal endothelial cells for SARS-CoV-2 virus. JCI Insight 6, 10.1172/jci.insight.148999.

Li, D., Edwards, R.J., Manne, K., Martinez, D.R., Schafer, A., Alam, S.M., Wiehe, K., Lu, X., Parks, R., Sutherland, L.L., et al. (2021). In vitro and in vivo functions of SARS-CoV-2 infection-enhancing and neutralizing antibodies. Cell 184, 4203–4219 e4232.

Liebschner, D., Afonine, P.V., Baker, M.L., Bunkoczi, G., Chen, V.B., Croll, T.I., Hintze, B., Hung, L.W., Jain, S., McCoy, A.J., et al. (2019). Macromolecular structure determination using X-rays, neutrons and electrons: recent developments in Phenix. Acta Crystallogr D Struct Biol 75, 861–877.

Lim, W.W., Mak, L., Leung, G.M., Cowling, B.J., and Peiris, M. (2021). Comparative immunogenicity of mRNA and inactivated vaccines against COVID-19. The Lancet Microbe 10.1016/S2666-5247(21)00177-4.

Liu, C., Ginn, H.M., Dejnirattisai, W., Supasa, P., Wang, B., Tuekprakhon, A., Nutalai, R., Zhou, D., Mentzer, A.J., Zhao, Y., et al. (2021a). Reduced neutralization of SARS-CoV-2 B.1.617 by vaccine and convalescent serum. Cell 184, 4220–4236 e4213.

Liu, L., Wang, P., Nair, M.S., Yu, J., Rapp, M., Wang, Q., Luo, Y., Chan, J.F., Sahi, V., Figueroa, A., et al. (2020). Potent neutralizing antibodies against multiple epitopes on SARS-CoV-2 spike. Nature 584, 450–456.

Liu, Y., Soh, W.T., Kishikawa, J.I., Hirose, M., Nakayama, E.E., Li, S., Sasai, M., Suzuki, T., Tada, A., Arakawa, A., et al. (2021b). An infectivity-enhancing site on the SARS-CoV-2 spike protein targeted by antibodies. Cell 184, 3452–3466 e3418.

Lopez Bernal, J., Andrews, N., Gower, C., Gallagher, E., Simmons, R., Thelwall, S., Stowe, J., Tessier, E., Groves, N., Dabrera, G., et al. (2021). Effectiveness of Covid-19 Vaccines against the B.1.617.2 (Delta) Variant. N Engl J Med 385, 585–594.

Madhi, S.A., Baillie, V., Cutland, C.L., Voysey, M., Koen, A.L., Fairlie, L., Padayachee, S.D., Dheda, K., Barnabas, S.L., Bhorat, Q.E., et al. (2021). Efficacy of the ChAdOx1 nCoV-19 Covid-19 Vaccine against the B.1.351 Variant. N Engl J Med 384, 1885–1898.

McCallum, M., Bassi, J., De Marco, A., Chen, A., Walls, A.C., Di Iulio, J., Tortorici, M.A., Navarro, M.J., Silacci-Fregni, C., Saliba, C., et al. (2021). SARS-CoV-2 immune evasion by the B.1.427/B.1.429 variant of concern. Science 373, 648–654.

Muik, A., Wallisch, A.K., Sanger, B., Swanson, K.A., Muhl, J., Chen, W., Cai, H., Maurus, D., Sarkar, R., Tureci, O., et al. (2021). Neutralization of SARS-CoV-2 lineage B.1.1.7 pseudovirus by BNT162b2 vaccine-elicited human sera. Science 371, 1152–1153.

Nimmerjahn, F., and Ravetch, J.V. (2008). Fcγ receptors as regulators of immune responses. Nat Rev Immunol 8, 34–47.

Pettersen, E.F., Goddard, T.D., Huang, C.C., Couch, G.S., Greenblatt, D.M., Meng, E.C., and Ferrin, T.E. (2004). UCSF Chimera--a visualization system for exploratory research and analysis. J Comput Chem 25, 1605–1612.

Planas, D., Bruel, T., Grzelak, L., Guivel-Benhassine, F., Staropoli, I., Porrot, F., Planchais, C., Buchrieser, J., Rajah, M.M., Bishop, E., et al. (2021a). Sensitivity of infectious SARS-CoV-2 B.1.1.7 and B.1.351 variants to neutralizing antibodies. Nat Med 27, 917–924.

Planas, D., Veyer, D., Baidaliuk, A., Staropoli, I., Guivel-Benhassine, F., Rajah, M.M., Planchais, C., Porrot, F., Robillard, N., Puech, J., et al. (2021b). Reduced sensitivity of SARS-CoV-2 variant Delta to antibody neutralization. Nature 596, 276–280.

Polack, F.P., Thomas, S.J., Kitchin, N., Absalon, J., Gurtman, A., Lockhart, S., Perez, J.L., Perez Marc, G., Moreira, E.D., Zerbini, C., et al. (2020). Safety and Efficacy of the BNT162b2 mRNA Covid-19 Vaccine. N Engl J Med 383, 2603–2615.

Priem, B., van Leent, M.M.T., Teunissen, A.J.P., Sofias, A.M., Mourits, V.P., Willemsen, L., Klein, E.D., Oosterwijk, R.S., Meerwaldt, A.E., Munitz, J., et al. (2020). Trained Immunity-Promoting Nanobiologic Therapy Suppresses Tumor Growth and Potentiates Checkpoint Inhibition. Cell 183, 786–801 e719.

Punjani, A., Rubinstein, J.L., Fleet, D.J., and Brubaker, M.A. (2017). cryoSPARC: algorithms for rapid unsupervised cryo-EM structure determination. Nat Methods 14, 290–296.

Robbiani, D.F., Gaebler, C., Muecksch, F., Lorenzi, J.C.C., Wang, Z., Cho, A., Agudelo, M., Barnes, C.O., Gazumyan, A., Finkin, S., et al. (2020). Convergent antibody responses to SARS-CoV-2 in convalescent individuals. Nature 584, 437–442.

Saito, F., Hirayasu, K., Satoh, T., Wang, C.W., Lusingu, J., Arimori, T., Shida, K., Palacpac, N.M.Q., Itagaki, S., Iwanaga, S., et al. (2017). Immune evasion of Plasmodium falciparum by RIFIN via inhibitory receptors. Nature 552, 101–105.

Schafer, A., Muecksch, F., Lorenzi, J.C.C., Leist, S.R., Cipolla, M., Bournazos, S., Schmidt, F., Maison, R.M., Gazumyan, A., Martinez, D.R., et al. (2021). Antibody potency, effector function, and combinations in protection and therapy for SARS-CoV-2 infection in vivo. J Exp Med 218.

Shrotri, M., Navaratnam, A.M.D., Nguyen, V., Byrne, T., Geismar, C., Fragaszy, E., Beale, S., Fong, W.L.E., Patel, P., Kovar, J., et al. (2021). Spike-antibody waning after second dose of BNT162b2 or ChAdOx1. The Lancet 398, 385–387.

Soh, W.T., Liu, Y., Nakayama, E.E., Ono, C., Torii, S., Nakagami, H., Matsuura, Y., Shioda, T., and Arase, H. (2020). The N-terminal domain of spike glycoprotein mediates SARS-CoV-2 infection by associating with L-SIGN and DC-SIGN. bioRxiv 10.1101/2020.11.05.369264.

Suryadevara, N., Shrihari, S., Gilchuk, P., VanBlargan, L.A., Binshtein, E., Zost, S.J., Nargi, R.S., Sutton, R.E., Winkler, E.S., Chen, E.C., et al. (2021). Neutralizing and protective human monoclonal antibodies recognizing the N-terminal domain of the SARS-CoV-2 spike protein. Cell 184, 2316–2331 e2315.

Tegally, H., Wilkinson, E., Giovanetti, M., Iranzadeh, A., Fonseca, V., Giandhari, J., Doolabh, D., Pillay, S., San, E.J., Msomi, N., et al. (2021). Detection of a SARS-CoV-2 variant of concern in South Africa. Nature 592, 438–443.

Thepaut, M., Luczkowiak, J., Vives, C., Labiod, N., Bally, I., Lasala, F., Grimoire, Y., Fenel, D., Sattin, S., Thielens, N., et al. (2021). DC/L-SIGN recognition of spike glycoprotein promotes SARS-CoV-2 trans-infection and can be inhibited by a glycomimetic antagonist. PLoS Pathog 17, e1009576.

Voss, W.N., Hou, Y.J., Johnson, N.V., Delidakis, G., Kim, J.E., Javanmardi, K., Horton, A.P., Bartzoka, F., Paresi, C.J., Tanno, Y., et al. (2021). Prevalent, protective, and convergent IgG recognition of SARS-CoV-2 non-RBD spike epitopes. Science 372, 1108–1112.

Wang, P., Nair, M.S., Liu, L., Iketani, S., Luo, Y., Guo, Y., Wang, M., Yu, J., Zhang, B., Kwong, P.D., et al. (2021a). Antibody resistance of SARS-CoV-2 variants B.1.351 and B.1.1.7. Nature 593, 130–135.

Wang, Z., Schmidt, F., Weisblum, Y., Muecksch, F., Barnes, C.O., Finkin, S., Schaefer-Babajew, D., Cipolla, M., Gaebler, C., Lieberman, J.A., et al. (2021b). mRNA vaccine-elicited antibodies to SARS-CoV-2 and circulating variants. Nature 592, 616–622.

Weisblum, Y., Schmidt, F., Zhang, F., DaSilva, J., Poston, D., Lorenzi, J.C., Muecksch, F., Rutkowska, M., Hoffmann, H.H., Michailidis, E., et al. (2020). Escape from neutralizing antibodies by SARS-CoV-2 spike protein variants. Elife 9, e61312.

Widge, A.T., Rouphael, N.G., Jackson, L.A., Anderson, E.J., Roberts, P.C., Makhene, M., Chappell, J.D., Denison, M.R., Stevens, L.J., Pruijssers, A.J., et al. (2021). Durability of Responses after SARS-CoV-2 mRNA-1273 Vaccination. N Engl J Med 384, 80–82.

Williams, C.J., Headd, J.J., Moriarty, N.W., Prisant, M.G., Videau, L.L., Deis, L.N., Verma, V., Keedy, D.A., Hintze, B.J., Chen, V.B., et al. (2018). MolProbity: More and better reference data for improved all-atom structure validation. Protein Sci 27, 293–315.

Winkler, E.S., Gilchuk, P., Yu, J., Bailey, A.L., Chen, R.E., Chong, Z., Zost, S.J., Jang, H., Huang, Y., Allen, J.D., et al. (2021). Human neutralizing antibodies against SARS-CoV-2 require intact Fc effector functions for optimal therapeutic protection. Cell 184, 1804–1820 e1816.

Xu, C., Wang, Y., Liu, C., Zhang, C., Han, W., Hong, X., Wang, Y., Hong, Q., Wang, S., Zhao, Q., et al. (2021). Conformational dynamics of SARS-CoV-2 trimeric spike glycoprotein in complex with receptor ACE2 revealed by cryo-EM. Sci Adv 7, eabe5575.

Yurkovetskiy, L., Wang, X., Pascal, K.E., Tomkins-Tinch, C., Nyalile, T.P., Wang, Y., Baum, A., Diehl, W.E., Dauphin, A., Carbone, C., et al. (2020). Structural and Functional Analysis of the D614G SARS-CoV-2 Spike Protein Variant. Cell 183, 739–751 e738.

Zost, S.J., Gilchuk, P., Chen, R.E., Case, J.B., Reidy, J.X., Trivette, A., Nargi, R.S., Sutton, R.E., Suryadevara, N., Chen, E.C., et al. (2020). Rapid isolation and profiling of a diverse panel of human monoclonal antibodies targeting the SARS-CoV-2 spike protein. Nat Med 26, 1422–1427.

